# Epigenome priming dictates transcription response and white matter fate upon perinatal inflammation

**DOI:** 10.1101/411702

**Authors:** Anne-Laure Schang, Juliette van Steenwinckel, Julia Lipecki, Charlotte Rich-Griffin, Kate Woolley-Allen, Nigel Dyer, Tifenn Le Charpentier, Patrick Schäfer, Bobbi Fleiss, Sascha Ott, Délara SabéRan-Djoneidi, Valérie Mezger, Pierre Gressens

**Author notes:** Equal contribution to the work (Juliette van Steenwinckel, Julia Lipecki). Equal contribution to the direction of the work (Sascha Ott, Délara SabéRan-Djoneidi, Valérie Mezger, Pierre Gressens). INSERM, UMR1153, Epidemiology and Biostatistics Sorbonne Paris Cité Center (CRESS), HERA team. Université de Paris, Faculté de Santé, Faculté de Pharmacie de Paris - 4 avenue de l’Observatoire – 75006 PARIS, France. School of Health and Biomedical Sciences, RMIT University, Bundoora, VIC, Australia. **Author’s contributions** ALS performed bioinformatics microarray and ATAC-seq dataset analyses, as well as wet experiments, including the MACS-isolation of O4+ OPCs and RT-qPCR experiments, in association with JvS (experimental design, mouse treatment and cell sorting) and TLC (cell culture, cytokine quantification by luminex). BF contributed to the design of the project, and brought very helpful suggestions along the experimental process and writing of manuscript. SO, DSD, PG and VM codirected the work. SO supervised JL, CRG, KWA and ND in their input ATAC-seq analyses and developed bioinformatic tools for testing for enrichment of paired motifs of transcription binding sites, and also proposed and performed the bioinformatic comparison between the epigenomic landscapes of OPCs and inflamed HAECs. PS co-supervised JL and CRG. DSD played a major role in suggesting the use of ATAC-Seq, animated the collaboration with SO, and performed the MACs-isolation of OPCs and ATAC-Seq data experiments with ALS whom she co-supervised with VM and PG. PG scientifically co-directed with VM the project. In particular, he and BF were at the origin of the MACS-isolation of the OPCs and produced OPC transcriptomic datasets and initiated their analyses. VM drove the study by proposing to analyze and intersect epigenomic data and transcriptomic data, for which she supervised DSD and co-supervised ALS; she wrote the paper with input from ALS, BF, SO, DSD, and PG. **Declaration of interests** The authors declare no competing interests.

## Abstract

Inflammatory insults accompanying prematurity provokes diffuse white matter injury (DWMI) which is associated with increased risk of neurodevelopmental disorders: pre-term infants have a 10 to 18-fold increased risk of developing autism spectrum disorders, compared to term infants. DWMI is due to maturation arrest in oligodendrocyte precursor cells (OPCs). Using integrated genome-wide approaches in a validated mouse perinatal model of DWMI, induced by systemic- and neuro-inflammation based on repeated interleukin-1B administrations, we show that neuroinflammation induces limited epigenomic disturbances in OPCs. In contrast, we unravel marked transcriptomic alterations of genes of the immune/inflammatory pathways, which are expressed in unstressed OPCs and physiologically downregulated along OPC maturation. Consistently, we observe that transcription factors of the inflammatory pathways occupy DNA both in unstressed and inflamed OPCs. Thus, rather than altering genome-wide chromatin accessibility, neuroinflammation takes advantage of open chromatin regions and deeply counteracts the stage-dependent downregulation of these active transcriptional programs. Therefore, our study opens new avenues for the future development of targeted approaches to protect preterm brains.

**Highlights:**

∘ Limited epigenomic impact of inflammation on OPC maturation blockade
∘ Major transcriptomic disturbances take advantage of a primed epigenetic landscape
∘ Proinflammatory genes are active in OPCs and downregulated upon maturation
∘ Neuroinflammation counteracts both this downregulation and maturation in OPCs

## INTRODUCTION

Premature birth, namely birth before 37 of 40 completed weeks, occurs in 8-13 % of all births worldwide and is the commonest cause of death and disability in children under 5 years of age (Delobel-Ayoub et al., 2009). Life-long morbidity is predominantly due to neurological damage, which altogether includes an array of effects, collectively called “encephalopathy of prematurity” (Volpe, 2009). Almost 10% of infants born before 33 weeks develop cerebral palsy and approximately 35% have persistent cognitive and neuropsychiatric deficits, including autism spectrum disorders and attention deficit/hyperactivity disorder (Bokobza et al., 2019). Although the most severe problems stem from extreme prematurity, even slight reductions in gestational length have significant adverse effects. One of the hallmarks of encephalopathy of prematurity is diffuse white matter injury (DWMI), which is considered a key target for neuroprotection and the prevention of long-lasting handicap. DWMI is due to oligodendrocyte maturation arrest, leading to hypomyelination and ultimately to defects in grey matter connectivity (Delobel-Ayoub et al., 2009; Ball et al., 2012; Billiards et al., 2008). In that context, neuroinflammation is a leading cause of encephalopathy of prematurity, serving as a central mediator of oligodendrocyte maturation defects and hypomyelination (Leviton and Gressens, 2007; Hagberg et al., 2015).

We have previously validated a mouse model of encephalopathy of prematurity that recapitulates arrest in oligodendrocyte maturation, hypomyelination, cognitive deficits and neuroinflammation, as seen clinically (Favrais et al., 2011; Krishnan et al., 2017; Shiow et al., 2017; Rangon et al., 2018). In this model, the common exposure of preterm-born infants to systemic and central inflammation (neuroinflammation) that is known to drive encephalopathy is mimicked by intraperitoneal (i.p.) administration of interleukin 1B (IL1B) from postnatal days 1 to 5 (P1-P5). Because in clinical conditions and diverse models of DWMI, males are more severely affected than females (Hagberg et al., 2015), the studies on oligodendrocyte precursor cells (OPCs) - and the present study - are performed in male animals. The developmental window (P1-P5) is equivalent to the high-risk window for encephalopathy of prematurity in infants, [23-32]-week gestational age. This paradigm of DWMI causes long-term myelination defects (Favrais et al., 2011), but the delicately choreographed programs that control OPC maturation, which are disturbed in this model - creating a cell fate issue - are not understood and remain to be enlightened.

Here, using a purified population of premyelinating OPCs, from this animal model, we have explored, through integrated genome-wide approaches, the contribution of epigenomic and transcriptomic disturbances to the OPC dysmaturation. We show that at P5 a limited number of chromatin regions are perturbed. We also find that the genes, whose expression levels are the most significantly altered by neuroinflammation, are involved in the immune system and inflammatory response. These genes, which include cytokines and chemokines, are unexpectedly also expressed by OPCs in normal conditions, in a developmentally regulated manner: their expression is higher at early stages and downregulated over the course of their maturation. The stage-dependent downregulation of these genes is perturbed in OPCs isolated from our inflammatory model of encephalopathy, which provokes a marked upregulation of their expression – likely participating to the failure of these cells to mature correctly.

These major transcriptomic disturbances surprisingly occur in genes which exhibit no overt changes in chromatin accessibility at P5. Indeed, our evidence suggests that neuroinflammation takes advantage of transcriptional programs that are active at the time of exposure – namely open chromatin regions - and disturbs their developmental regulation.

We have therefore unraveled a mechanism by which neuroinflammation acts on OPCs and could participate in maturation arrest in a model mimicking perinatal inflammation in preterm born infant. Indeed, the unexpected expression of numerous inflammatory genes by OPCs during their normal stage-dependent maturation likely paves the way for intricate interference between the response to neuroinflammatory insults and the white matter developmental program, with important implications for therapeutic strategies.

## RESULTS

### Purity and quality of the O4+-purified cell populations used for exploration of the epigenome and transcriptome

In our previous studies, we have demonstrated that oligodendrocyte maturation arrest is a hallmark of the neuropathy caused by neuroinflammation that is triggered by intraperitoneal IL1B administration (IP). As outlined above and in Figure 1, IL1B was administered between P1 and P5 (versus PBS as a control; Figure 1; (Favrais et al., 2011; Krishnan et al., 2017; Shiow et al., 2017; Rangon et al., 2018); this period mimics chronic exposure to systemic neuroinflammatory mediators from 23-32 weeks of gestation in the infant. Using magnetic-activated cell sorting (MACS), we isolated at P5 the premyelinating OPC population (O4+ OPCs) from male cortices in each condition (Figure 1). O4 is considered a pan-marker of late OPCs in humans and mice (Volpe, 2009; Favrais et al., 2011). We evaluated the purity of the O4+ cell population by a panel of diverse approaches at each step during the course of our study (Figure S1). Notably, O4+ OPC cells were directly isolated from the pup cortices, and immediately and directly processed without culturing. By performing RT-qPCR analyses, we demonstrated that this population, expressed *Myelin binding protein* (*Mbp*) mRNAs, a marker of myelinating oligodendrocytes, whose transcription starts in pre-myelinating oligodendrocytes, contrary to microglial (CD11B+) MACS-isolated fractions (Figure S1A). Conversely, we found that the levels of the microglia marker mRNAs, CD11B (*Itgam, Integrin alpha M* gene) were almost undetectable in this O4+ cell population. This population therefore corresponds to premyelinating OPCs. In independent RT-qPCR experiments, as expected based on previous studies using this model (Favrais et al., 2011), we observed that the expression of *Id2*, encoding a transcriptional inhibitor of oligodendrocyte differentiation, was increased in O4+ OPCs in response to neuroinflammation, whereas that of myelination-associated genes was slightly, but reproducibly, reduced (Figure S2A).

**Figure 1.**
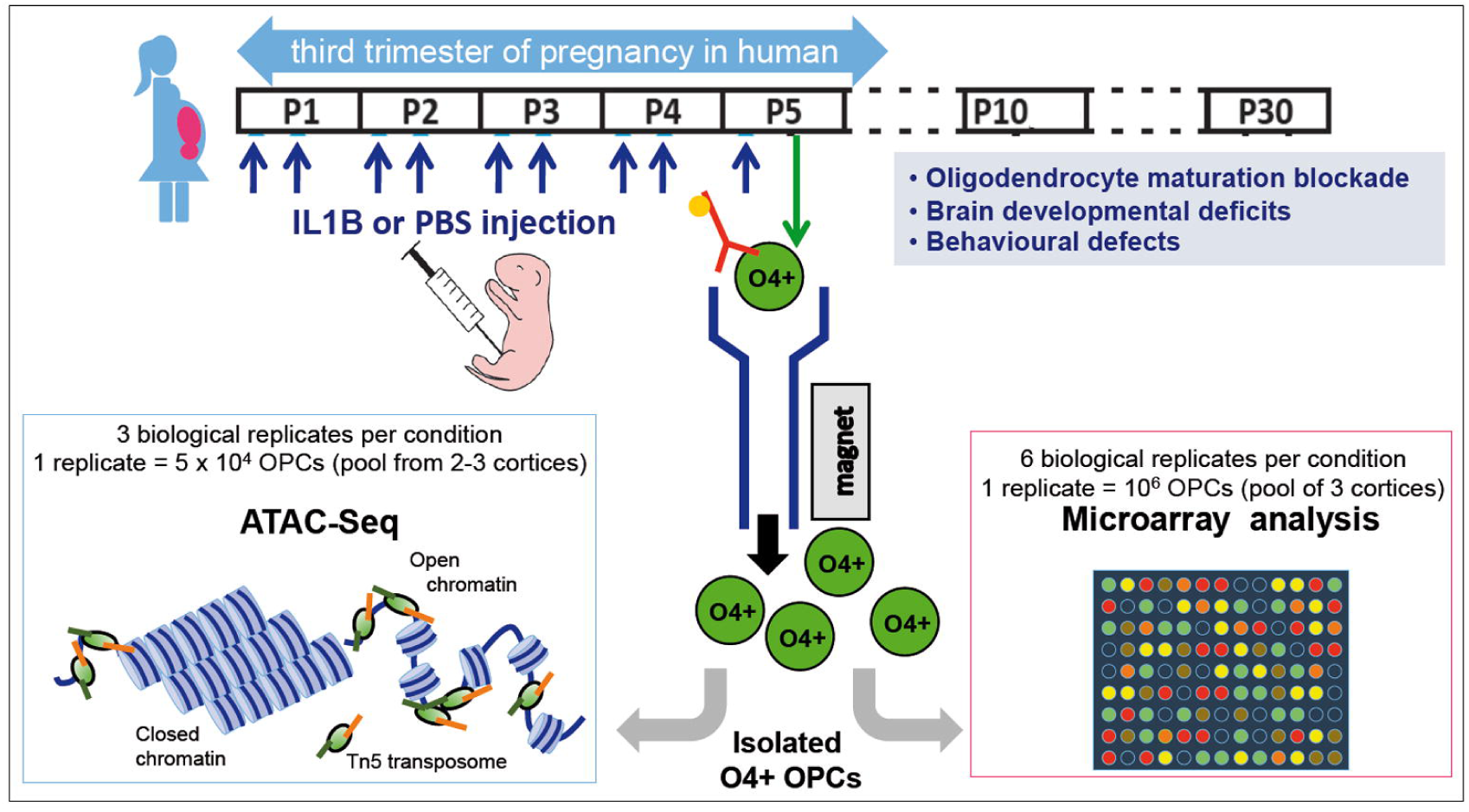
Experimental strategy. (see also Figure S1). Previously validated mouse model of encephalopathy of prematurity in which we mimic the systemic and neuroinflammatory insults as undergone by human infants from approximately [23-32]-week gestational age equivalent. Neuroinflammation is induced *via* i.p. IL1B from postnatal days 1 to 5, and this leads to OPC maturation blockade, defective myelination, and behavioral anomalies as seen clinically (Favrais et al., 2011). O4+ OPCs (green circles) were isolated from P5 pup cortices by MACS and genome-wide chromatin accessibility was explored by ATAC-Seq (lower, left panel): the enzymatic severing of the DNA by the transposome (Tn5 transposase, loaded with adapters *in vitro*; green and orange) allows « tagmentation » of DNA template, to fragments tagged with adapters. In parallel (lower, right panel), comparative transcriptomic analysis was performed using Agilent mouse whole genome microarray.

Overall, these data show that the cell population isolated by our MACS-protocol is predominantly enriched with O4+ OPCs and exhibits, as expected, hallmarks of maturation arrest in response to neuroinflammation induced by intraperitoneal administration of IL1B.

### The epigenome of OPCs is globally preserved in response to systemic IL1B exposure

We first investigated the impact of neuroinflammation the integrity of the chromatin landscape in OPCs using ATAC-Seq (Buenrostro et al., 2013; *Assay for Transposase-Accessible Chromatin with high-throughput sequencing*; Figure 1). Using the bioinformatics workflow described in Figure S2B (see Material and Methods) including the MACS2 (Zhang et al., 2008) and EdgeR (Robinson et al., 2010) software tools, we obtained an average of 72 million Tn5 transposase-integrated mapped reads per sample, representing a total of 213,246 statistically significant peaks (MACS2; FDR < 0.05; Table S1 and S2; Dataset S1). Analysis of the insert size distributions showed the expected nucleosome-induced pattern and 10.4bp periodicity with good consistency across samples, an indication of high data quality (Figure S2C; Buenrostro et al., 2015).

The number of reads, which reflects chromatin accessibility, was determined for each sample in the 213,246 peaks. We performed principal component analysis (PCA) on our samples and observed that principal component 1 (PC1) accounted for 42% of the variance and separated samples from control and neuroinflammation-exposed OPCs (Figure 2A). We obtained similar results using the EdgeR MDS function (Figure S3A). This shows that a large proportion of the variance in this dataset can be explained by the exposure to neuroinflammation. Among the 213,246 significant peaks, only 524 regions were open or closed in response to IL1B (FDR < 0.05, Figure 2B; Figure S3B; Table S3). The majority of peaks with differential chromatin accessibility was more open in IL1B, compared to PBS conditions (Figure 2B,C). The extent of changes in chromatin accessibility was small, with fold-changes of most peaks only mildly deviating from sample-to-sample variability (Figure 2B). The median fold-change of differentially open regions was 2.03-fold.

**Figure 2.**
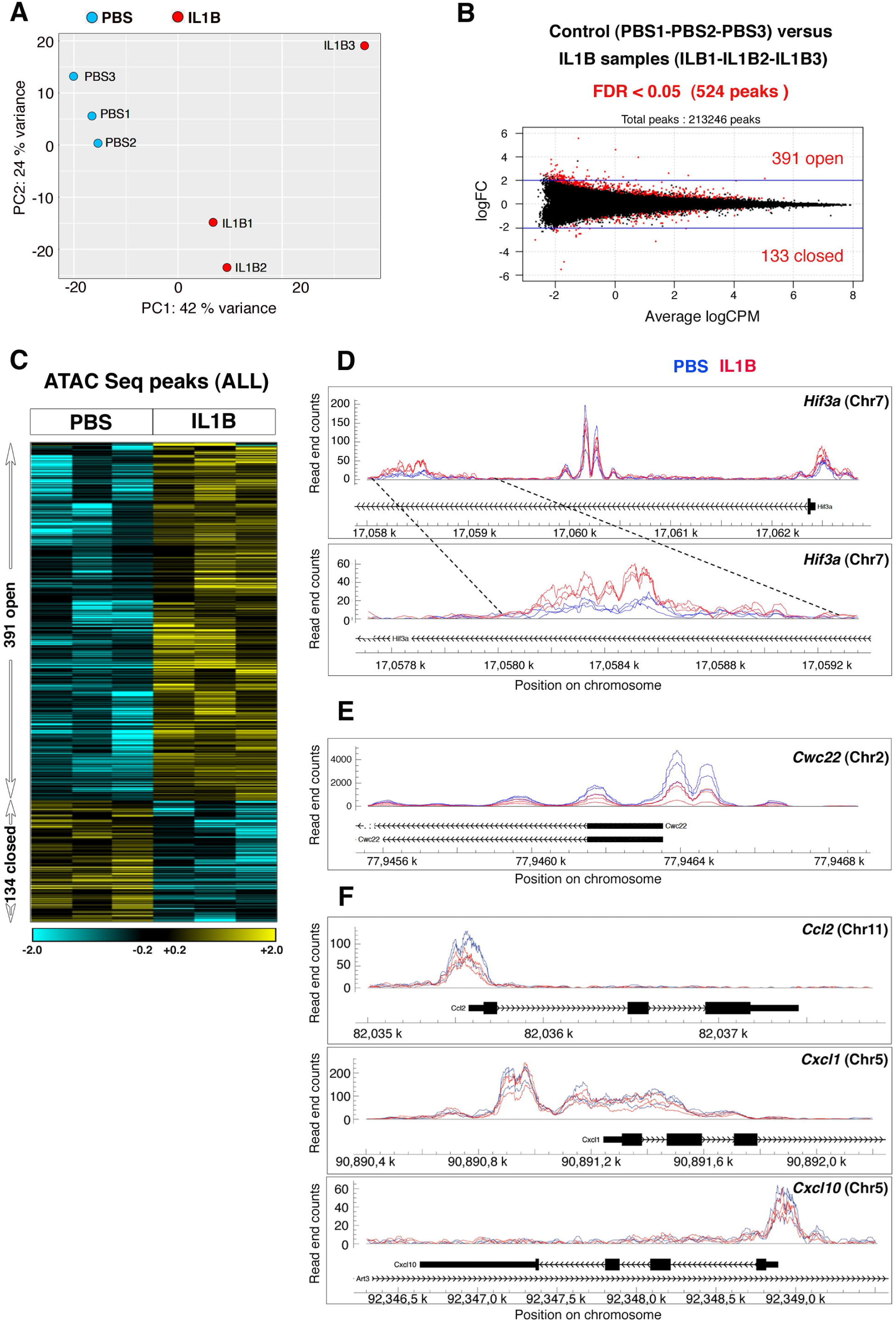
The epigenome of OPCs is globally preserved after IL1B treatment (ATAC-Seq analysis). (see Figure S1, S2 and S3) (**A**) PCA analysis of log normalized read counts falling within ATAC-seq chromatin peaks from OPCs from control (PBS) or neuroinflammation-exposed (IL1B injections); total number of peaks, 213,246. (**B**) Scatter plot representing the dispersion (fold change) of peaks in relation to the number of tn5 cut*<u>s</u>* per million (logCPM), for each individual analyzed peak across 3 PBS samples and 3 IL1B samples. The 524 peaks showing significant differences between IL1B and PBS conditions are indicated in red, with 391 peaks corresponding to increased and 133 peaks to decreased chromatin accessibility, with FDR < 0.05. (**C**) Heat map visualization of the 524 opening (yellow) or closing (blue) peaks in control (PBS) and neuroinflammation-exposed (IL1B injected) conditions. (**D**) Examples of peaks showing increased or unmodified chromatin accessibility: (Upper panel) the peaks located at the most downstream position in the *Hif3a gene* (*Hypoxia inducible factor 3 alpha subunit*) show increased chromatin accessibility in neuroinflammation context) Magnification of this region is illustrated in the lower panel). In contrast, peaks located in the middle of the *Hif3a* gene show no significant changes in chromatin accessibility (Upper panel). (**E**) Example of peak showing reduced chromatin accessibility: peaks within the *Cwc22* gene (encoding the spliceosome-associated protein 22). (**F**) Example of peaks in genes encoding cytokines and chemokines, showing no statistically relevant modification of chromatin accessibility.

We next annotated the 213,246 peaks using HOMER annotatePeaks function (Heinz et al., 2010). Peaks were mostly distributed in intergenic and intronic regions and were enriched in gene regulatory regions and gene bodies (Figure S3C). A similar distribution was observed for the 524 differential peaks (Figure S3C). Representative peaks showing increased, reduced, or unchanged chromatin accessibility are illustrated in Figure 2D-F. We assigned differential peaks to genes, likely affected by chromatin alteration, based on proximity to their transcription start site (TSS), and we performed gene ontology (GO) analyses, using David6.8 (Huang et al., 2009a; Huang et al., 2009b) for the 485 corresponding genes, of which 344 were assigned a GO-term. The top ten GO-terms ranked according to their FDR, belong to pathways relevant for neural development (regulation of cell migration, cell adhesion, and neuron projection development), but none of them showed FDR < 0.05 (Figure S3D; Table S4).

Our results thus reveal that only 0.25% of open chromatin regions show differential chromatin accessibility in P5 OPCs that are isolated from a model of inflammation-driven encephalopathy, showing that the chromatin landscape is globally preserved, in terms of accessibility,

### Major transcriptomic impact of neuroinflammation on the immune/inflammatory pathway in O4+ OPCs

We analyzed gene expression in isolated O4+ OPCs using microarray analysis. We compared six independent samples of O4+ OPCs from IL1B-exposed mice to six independent samples from PBS-treated (control) mice. Induced-neuroinflammation mainly triggered upregulation of gene expression: 1,266 genes were up-regulated and 454 downregulated, which corresponded to 1,872 and 699 probes, respectively (FC +/-1.5; FDR < 0.05; Figure 3A; Table S5). As expected based on previous work in this model (Favrais et al., 2011) and from our validation experiments by RT-qPCR for O4+ OPC isolation (Figure S2A), the analysis of our microarray data revealed that the expression of genes associated with myelination (*Mbp, Mog, Mag, CNPase, Plp1*; which are still lowly expressed at P5 in normal conditions) were downregulated by neuroinflammation, whereas that of *Id2* was upregulated (Favrais et al., 2011); Figure S4A). These results likely reflected OPC maturation blockade (Favrais et al., 2011). Strikingly our GO analysis of these upregulated genes pinpointed the immune system and inflammatory response in the top 5 most statistically significant pathways (DAVID 6.8; Figure 3B; Table S6). The analysis of downregulated genes indicated that these belonged to pathways linked to development, but with lower statistical relevance (Figure S4B). Interestingly, the vast majority of the genes belonging to the immune system and inflammatory response pathways and whose expression was significantly altered, appeared to be upregulated (220/262); Figure 3C). Using RT-qPCR on independent samples, we confirmed the induction of the expression of known players of these pathways: cytokines, chemokines, interleukins and their receptors (Figure 3D).

**Figure 3.**
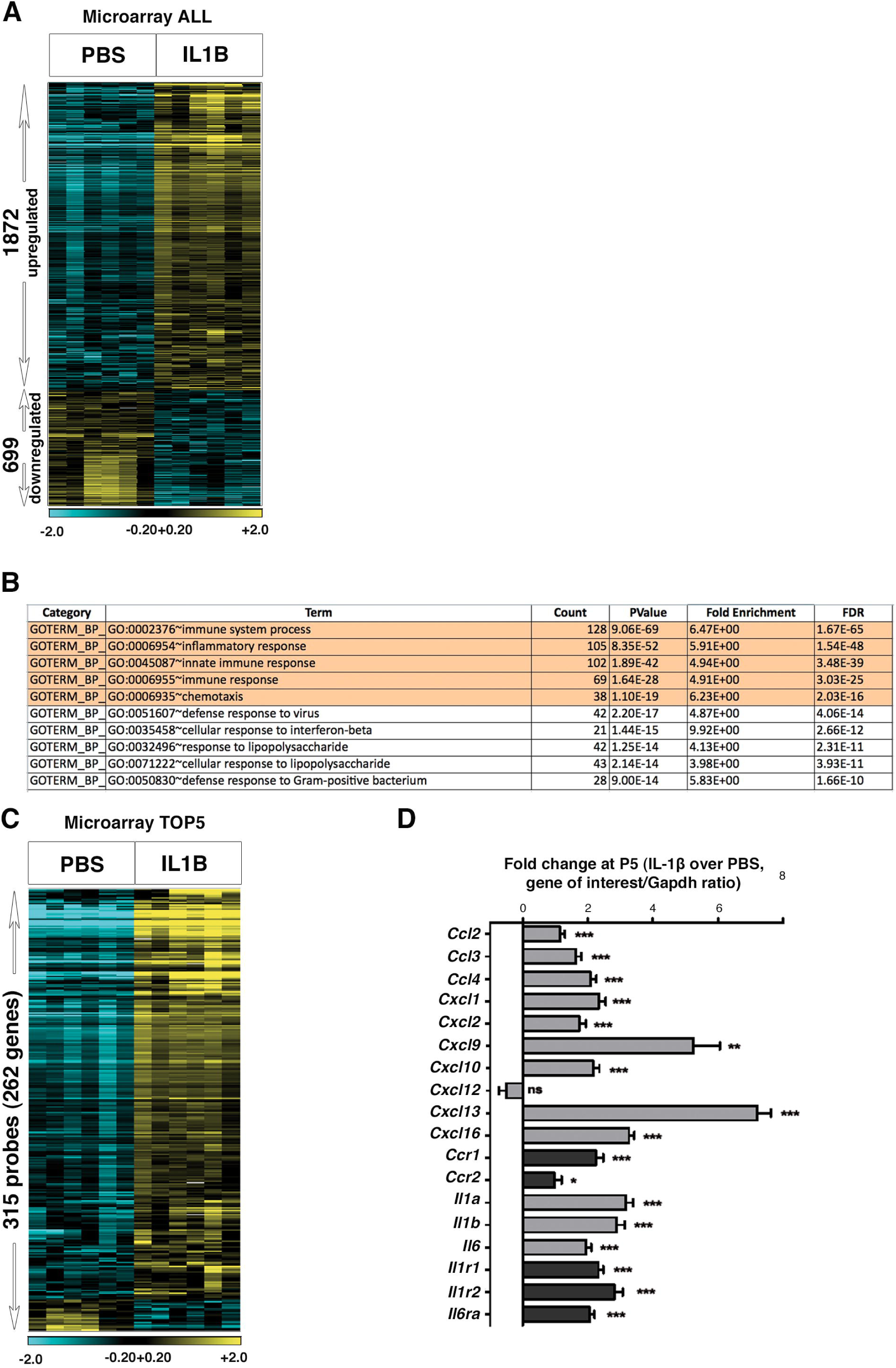
Major transcriptomic impact of neuroinflammation on the immune/inflammatory pathway in O4+ OPCs. (see Figures S1, S4A,B and Table S4). Microarray analysis comparing gene expression in isolated O4+ OPCs from six independent control (PBS) and six independent neuroinflammation-exposed cortices (IL1B; Figure 1). (**A**) ***Heat map of genes with altered expression upon IL1B exposure***. A fold-change (FC) threshold of +/-1.5 was chosen, with FDR < 0.05. Total number of probes corresponding to IL1B-induced disturbances in gene expression: 2571, corresponding to 1719 genes. Total number of probes corresponding to the 1266 upregulated genes: 1872. Total number of probes corresponding to the 454 downregulated genes: 699. Heatmap color scale: log2 [-2.0;+2.0]. (**B**) ***The top5 GO-terms of up-regulated genes corresponds to the immune system and inflammatory response pathways***. GO-terms as a table with the number of upregulated genes per GO-term, p-values, and FDR. (**C**) ***Heat map of the genes belonging to the top5 GO-terms***, 315 probes corresponding to 262 genes. Heatmap color scale: log2 [-2.0; +2.0]. (**D**) ***Validation of the alteration in gene expression for members of the GO-term “immune system and inflammatory response pathways” in O4+ OPCs using RT-qPCR***. Number of independent experiments: n=8 for all genes, except for *Cxcl9, Cxcl10*, and *Il1r1* (n=7).*, p < 0.05; **, p < 0.01; ***, p < 0.001. Grey bars ligands; black bars: receptors.

### OPCs intrinsically produce cytokine and chemokines in response to IL1B

We confirmed that cytokine and chemokine genes were upregulated by neuroinflammation in O4+ OPCs *per se*, and excluded the possibility that the upregulation of immune and inflammatory genes was due to contamination of O4+ OPCs with microglia, using different approaches. In this model of neuroinflammation, we have previously published microarray analyses of the transcriptomic profiles in microglia (CD11B+ MACS-isolated MG cells; (Krishnan et al., 2017; Van Steenwinckel et al., 2019). These CD11B+ cells were obtained from the same animals as the O4+ OPC populations from this study, by sequential MACS-based isolation of O4+ OPCs and CD11B+ microglia cells. We compared the microarray gene expression profiles in the previously assessed CD11B+ cells to the list of 262 genes found in OPCs and corresponding to inflammation and immune pathways (Figure 4A; Figure S1B; dataset from (Krishnan et al., 2017). O4+ OPCs and CD11B+ MG populations exhibited remarkable differences in gene expression profiles in response to neuroinflammation, both in the magnitude and direction of expression changes (Figure 4A; Figure S1B). Notably, the upregulation of these genes in microglia was greatest at P1 and their expression at P5 has already recovered, reaching basal levels comparable to that of PBS samples, while in OPCs their expression remained elevated (Krishnan et al., 2017; Figure 4B). Those data were confirmed by RT-qPCR in independent O4+ OPC and CD11B+ MG samples, and extended to astrocytes (GLAST+ MACS isolation; Figure 4B). Specifically, neither CD11B+ MG, nor GLAST+ astrocytes showed any increase in the expression of selected cytokines and chemokines at P5, in contrast to O4+ OPCs (illustrated here for *Ccl2, Cxcl1* and *Cxcl10*; Figure 4B). Altogether, these results indicate that the upregulation of immune and inflammatory pathway in O4+ OPCs in response to neuroinflammation in our microarray analyses at P5 cannot be attributed to contamination of OPCs by microglia, nor astrocytes.

**Figure 4.**
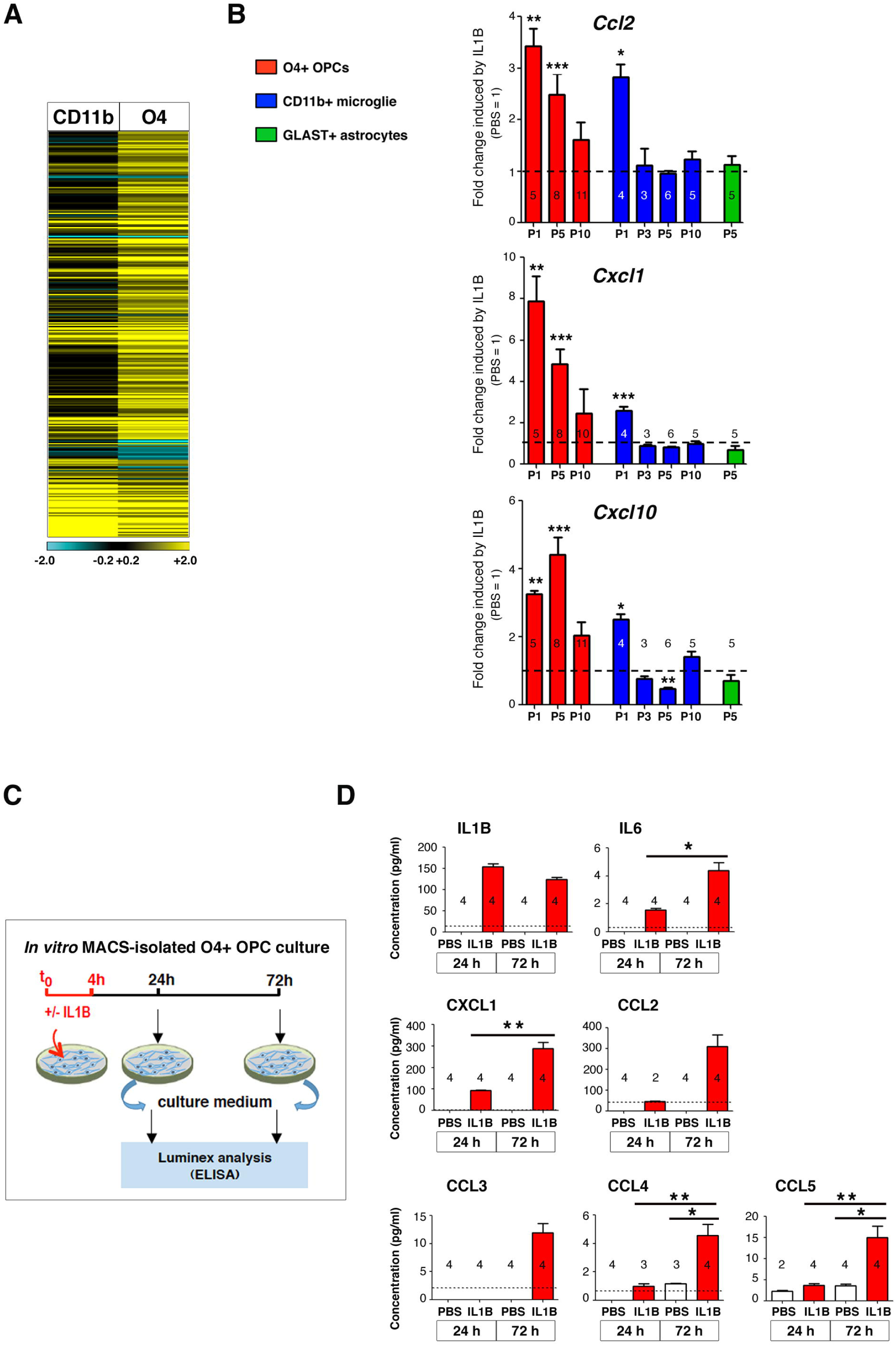
The production of cytokine and chemokine proteins by MACS-isolated O4+ OPCs is induced by neuroinflammation *in vivo* and *ex vivo*. (see Figures S1, S4C) (**A**) ***Isolated O4+ OPCs and isolated CD11B+ microglial cells exhibit distinct induction profiles of immune/inflammatory gene expression in response to neuroinflammation***. Heatmap comparison of the microarray data of the 262 upregulated genes of the immune/inflammatory pathway in isolated O4+ (described in Figure 3C) with the corresponding genes in CD11B+ microglial cells (Krishnan et al., 2017). Notably, these data where produced from cells originating from the same brains, at the same timepoint. Heatmap color scale of log2 [-2.0;+2.0]. (**B**) ***Unique signature for inflammatory gene expression in isolated O4+ OPCs, compared to CD11B+ microglia, and GLAST+ astrocytes***. Fold-change in the expression of genes of the immune and inflammatory pathways as detected by RT-qPCR analyses at different postnatal stages. mRNA levels are normalized to *Gapdh* for OPCs and astrocytes and *Rpl13* for microglia based on in-house reference gene testing. The numbers of independent experiments are indicated on each plot. *, p < 0.05; **, p < 0.01; ***, p < 0.001. (**C**) ***Experimental design for ex vivo OPC culture, inflammatory exposure (IL1B), and differentiation*** (see Material and Methods). (**D**) ***Protein detection and quantification*** (pG/mL) of the expression of interleukins (IL1B, IL6), cytokine C-C Motif Chemokine Ligand 2, 4, and 5 (CCL2, CCL3, CCL4, and CCL5) and chemokine C-X-C Motif Chemokine Ligand 2 (CXCL2) by Luminex. CTR, PBS exposure; IL1B, IL1B exposure. The dotted line represents the limit of detection for individual proteins in the assay. The numbers of independent experiments performed for each plot are indicated on each plot. *, p < 0.05; **, p < 0.01; ***, p< 0.001.

In addition, we also ruled out the possibility that a very low number of contaminating MG would be responsible for producing cytokine and chemokine transcripts, by investigating the chromatin accessibility of genes specific for OPCs or MG in ATAC-Seq analyses. We showed that *Olig2*, a pan-oligodendrocyte marker, which is not expressed by MG, displayed open chromatin structure in our O4+ OPC population, as indicated by the presence of ATAC-Seq peaks, but showed closed chromatin conformation in purified MG (data mining from Matcovitch-Natan et al. 2016; Figure S1C,D). Conversely, as expected, the MG-specific gene, *Itgam*, exhibited closed chromatin conformation in O4+ OPCs, in contrast to what was observed in MG (Figure S1C). So, if the cytokine and chemokine transcripts were produced by contaminant MG and not by OPCs, we should have seen a stronger ATAC-seq peak in MG than in OPCs at corresponding genes. This is not what we observed. When investigating the chromatin status of cytokine and chemokine genes in our MACS O4+ OPC population, we detected ATAC-Seq peaks with two different profiles as illustrated in Figure S1D: i) *Cxcl1*, that was found in an open chromatin state in the MACS-isolated O4+ OPCs population, but not in isolated MG. This indicates that this gene is in an open chromatin conformation in OPCs, whereas it is not in MG. This therefore implies that MG cannot be the major contributor to the synthesis of these molecules in the MACS-isolated O4+ OPC population; ii) *Ccl2* or *Cxcl10* were found in an open conformation in MACS-isolated O4+ OPCs, as well as in isolated MG, but, since *Itgam* was not detected as open in the MACS-isolated O4+ OPC population, it means that these peaks mainly correspond to the *Ccl2* or *Cxcl10* gene status that exists within MACS-isolated O4+ OPCs, and not within MG (Figure S1D; note that the peak profiles of *Cxcl1, Ccl2* or *Cxcl10* are also illustrated in Figure 2F, but, in that case, separately showing PBS1-3 and IL1B 1-3 samples).

We concluded that contamination of MACS-isolated O4+ OPCs by MG, if any, is only very minor in this study, as it is undetectable in our ATAC-Seq experiments. Such contamination therefore cannot account for the open chromatin status detected in cytokine or chemokine genes in the MACS-isolated O4+ OPC population.

To further verify the potential for production of inflammatory players by OPCs in response to inflammation, we isolated primary OPCs, grew them *in vitro* and treated these cultures with IL1B (Figure 4C). We detected significant induction of the production of seven cytokine and chemokine proteins *via* Luminex in the supernatant of IL1B-treated MACS-isolated primary O4+ OPCs, cultured for 24 or 72 hours (Figure 4D). We also confirmed the upregulation of these inflammatory and immune mediators at mRNA levels (Figure S4C).

In conclusion, these data strongly support the hypothesis that, at P5, O4+ OPCs are able to intrinsically synthesize inflammatory and immune pathway proteins in response to exposure to inflammatory stimuli.

### Major involvement of transcription factors of the immune and inflammatory pathways in O4+ OPCs

To globally examine the contribution of modifications of chromatin accessibility to the transcriptomic changes, we intersected our ATAC-Seq data (213,246 peaks, corresponding to 20,108 gene names) and our microarray data (limited to 25,294 genes with annotated ID). We found 16,883 genes in common between the two datasets, of which 1,333 genes showed altered expression after IL1B exposure and 404 peaks showed differential chromatin accessibility. By performing the intersection of the 1,333 differentially expressed genes with genes located nearby the differential 404 ATAC-Seq peaks, we identified 53 genes representing a statistically significant overlap between transcriptomic and epigenomic changes. (Figure 5A; p= 1.7 e^-4^; hypergeometric test; Table S7).

**Figure 5.**
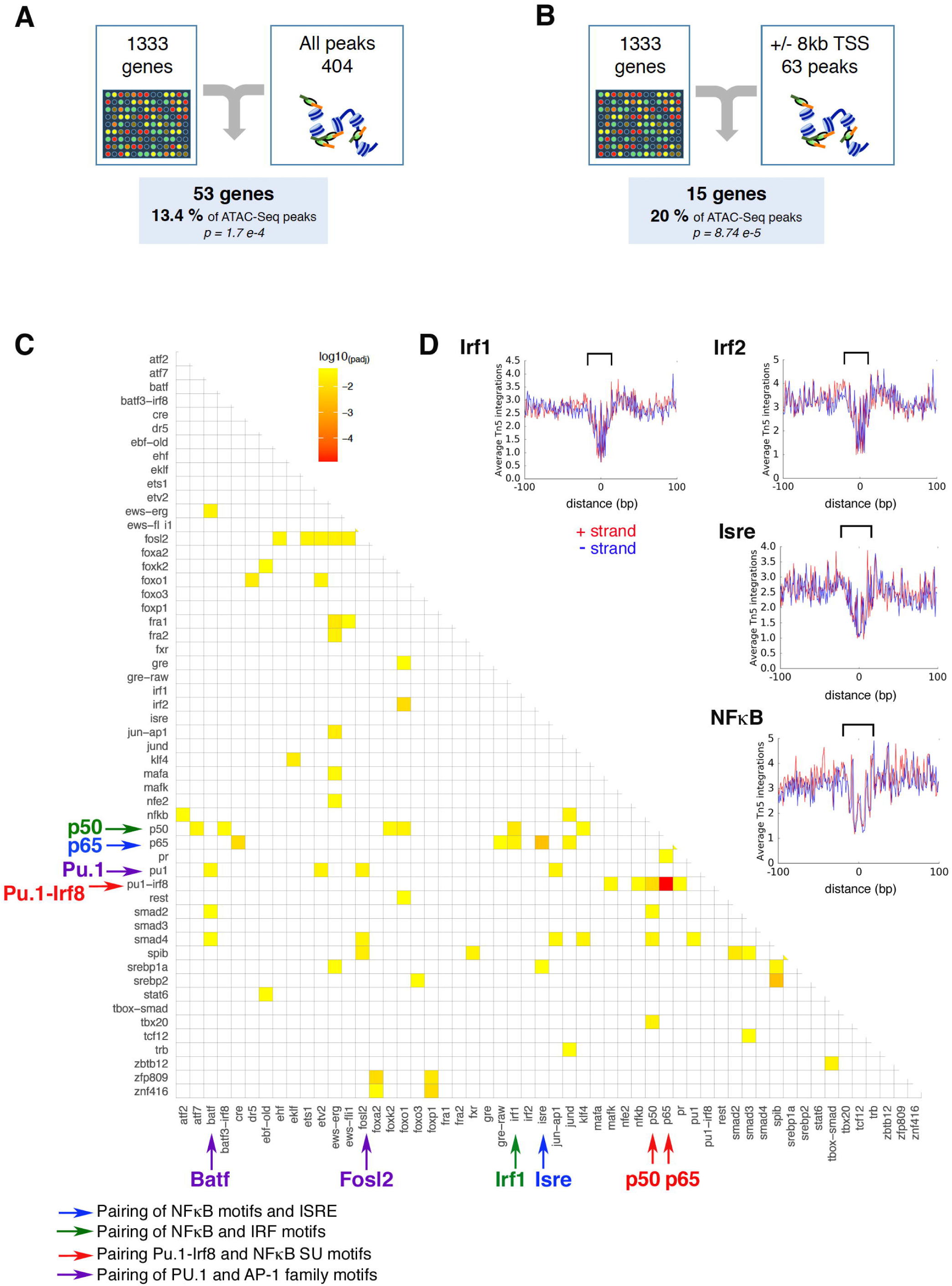
Major involvement of TFs of the immune and inflammatory pathways in O4+ OPCs. (see Figures S5 and S6) (**A** and **B**) ***Crosstabulation of ATAC-Seq and microarray data***. (**A**) Number of genes differentially expressed and associated with all differentially accessible peaks/regions in O4+ OPCs isolated from neuroinflammation-exposed pups. (**B**) Number of genes differentially expressed and associated with differentially accessible peaks/regions only located +/- 8 kb around their TSS in O4+ OPCs isolated from neuroinflammation-exposed pups. (**C**) ***Co-localization of pairs of TF binding motifs in ATAC-Seq peaks of upregulated genes***. Heat map of the paired motifs located in the 2319 ATAC-Seq peaks located within +/- 8 kb from the TSS of 1266 upregulated genes (corresponding to 886 different gene names), identified using paired motif analysis. Transcription factors (TF) names from MEME nomenclature on both axes. Statistically relevant co-occurrence of two different TFs is indicated by yellow to orange squares (multiple testing corrected log10 p-value). Examples of TF pairs are indicated IRF1-p50; *ISRE*-p65; PU.1-IRF8-p65; PU.1-iRF8-p50; PU.1-BATF; PU.1-FOSL2. Color code (TF names and arrows): red, pairing of PU.1-IRF8 with NFκB subunit binding motifs; purple, pairing of PU.1 with AP-1 family binding motifs; blue, pairing of NFΚB binding motifs with *Isre* (IFN-stimulated response element); green, pairing of NFΚB with IRF binding motifs. (**D**) Chromatin accessibility footprints at TF motifs. Examples of average footprint profiles at TF motifs that were plotted at base pair resolution, located within accessible chromatin *loci* in <u>both</u> PBS and IL1B conditions, adjacent to differentially regulated genes (ALL; down and upregulated) using the Wellington software (Piper et al., 2013). Typical footprint profiles coincide with the center of the canonical motif for a given TF (indicated by brackets). Footprints for occupancy of IRF1, IRF2, *Isre*, and NFκB binding motifs are illustrated. Footprints showing occupancy by IRF1, IRF2, *Isre*, and NFκB on corresponding binding motifs are illustrated. Red and blue curves: + and - strand, respectively.

In order to maximize the chance of attributing peaks to relevant genes, we reasoned that peaks corresponding to opening or closing of the chromatin and lying within +/- 8kb around a TSS were most likely to contribute to the regulation of the expression of the corresponding gene. Taking this into account, we identified 15 genes showing differential expression and exhibiting modifications of the chromatin landscape in their vicinity, which corresponded to 20 % of the ATAC-Seq peaks present in +/- 8 kb around the TSS (Figure 5B; p= 8.74 e^-5^; Table S8). Among these 15 genes, 10 were involved in the immune system and inflammatory response pathways: *Cd14, Cwc22* (Figure 2E), *Hmha1, Ifit3, March1, Nckap1l, Slfn2, Slc15a3, Tlr1, Tnfsf14, Tnfrsf12a* (Table S8). Interestingly, *Hif3a*, a gene recently identified in models of inflammation, was also included in this list (see discussion; Figure 2D).

In summary, the immune system and inflammatory response pathways were prominently represented in the lists of genes showing marked dysregulation of their expression upon exposure to neuroinflammation, and associated to two different chromatin behaviors: 1) a limited number of genes showed differential chromatin accessibility upon IL1B exposure (Figure 5A,B); 2) the majority of the genes annotated in the top 5 most enriched GO-terms displayed no major changes in chromatin accessibility. Indeed, in this case, the chromatin was already in an open conformation in the PBS samples (examples shown in Figure 2F).

### Identification of combinatorial transcription factor binding in immune system and inflammatory pathways

In the large majority of instances where neuroinflammation altered gene expression, we observed that the chromatin was already open and remained unchanged in response to neuroinflammation (examples shown in Figure 2F). As such, we investigated the putative involvement of transcriptional regulators as primary mediators of alteration in gene expression. We searched for enrichment in transcription factor (TF) binding sites (TFBS) using HOMER and known motifs in the ATAC-Seq peaks adjacent to differentially regulated genes (up and downregulated genes, termed “ALL”). We chose to focus on ATAC-Seq peaks within a distance of +/- 8 kb relative to the TSS of these genes. Motifs for members of the IRF (interferon-regulatory factor) family appeared at the top of the list with the strongest scoring results (Figure S5A). A list of similar motifs, including motifs for the NFκB family members, was found with comparable scoring results, as well as p-values, in the peaks adjacent to upregulated genes (termed “UP”; Figure S5B). Notably, the composite site PU.1-IRF8 and were enriched in “UP” genes (Figure S5B).

Because we suspected that these TFs might work together (Yanai et al., 2012), we investigated the occurrence of paired motifs in the peaks located in +/- 8kb regions around the TSS of differentially regulated genes, as described in Materials and Methods. The analysis of peaks corresponding to downregulated genes did not reveal any paired motif enrichment, compared to random occurrence in all peaks. In contrast, the analyses of ATAC-Seq peaks associated with upregulated genes revealed the existence of paired TFBS motifs, with marked involvement of TFBS from the IRF family, PU.1/SPI1, *Isre* (Interferon-Stimulated Response Element), NFκB, and AP-1 family (Figure 5C; Table S9).

To investigate whether the occurrence of these motifs corresponded to the binding of TFs, we tested the motif occupancy on DNA in PBS and IL1B conditions. For this, we used the Wellington algorithm (Piper et al., 2013), which is highly accurate in inferring protein (TF)-DNA interactions. We investigated the presence of footprints corresponding to occupied TFBS and their motif content, located within significant ATAC-Seq peaks, adjacent to differentially regulated genes. The average footprint profiles, produced for the top 6, high-ranked, HOMER motifs (Figure S5A,B) are illustrated for IRF1, IRF2, IRSE and NFκB in Figure 5D (using data from both conditions, PBS plus IL1B) and in Figure S6A (PBS, upper panels; IL1B, lower panels). The dip in the number of reads at the center of the sharp average profile (indicated by brackets) was strongly suggestive of effective TF binding (Figure 5D and S6A). In contrast, PU.1-IRF8 and PGR (Progesterone Receptor) average footprint exhibited sharp internal spikes, suggestive of transposase insertion bias (Green et al., 2012, Figure S6B). Interestingly, there was little difference in the average footprint profiles for all these TFs, when considering either PBS samples only or IL1B samples only, indicating that these TFs (or TFs binding similar motifs), might already bind DNA at the corresponding motifs in unstressed, physiological conditions (Figure S6A).

Altogether, these results indicate that key TFs involved in the immunity and inflammatory processes are bound in a significant number of open regions located in the vicinity of genes that are differentially upregulated by neuroinflammation. The binding of such TFs is coherent with the prominent upregulation of proinflammatory cytokine and chemokine genes, and other genes of the immune system and inflammatory pathways. In addition, because footprint profiles were similar in PBS and IL1B conditions, our data suggest that, at P5, these TFs might be positioned in these regions before exposure to neuroinflammation.

There was no evidence for footprints in peaks adjacent to downregulated genes (data not shown). Moreover, the search for de *novo* motifs in ATAC-Seq peaks near differentially expressed genes did not reveal statistically relevant motif associated with *bona fide* average footprints (data not shown).

### Constitutive expression of genes of the immune and inflammatory pathways at early stages of OPC maturation trajectory, in unstressed conditions

At this step of the study, we had shown that P5 isolated O4+ OPCs were able to induce the expression of genes belonging to the immune system and inflammatory response pathways, in response to neuroinflammatory challenge. We had also shown that these major alterations in gene expression occurred without major modifications in chromatin accessibility, but that key TFs controlling these pathways were bound to these open regions both in the control (PBS) and neuroinflammatory (IL1B) conditions. These observations suggest that the epigenomic profiles associated with neuroinflammation-induced genes exhibit “neuroinflammatory-like” patterns, prior to IL1B injections (*i*.*e*. open chromatin conformation). Therefore, we wondered if the transcription of these genes may already be activated at a physiological level in OPCs during normal development and further upregulated by exposure to neuroinflammation.

To assess this hypothesis, we MACS-isolated O4+ OPCs at P3, P5, and P10 from our model of preterm neuroinflammatory injury, and indeed, we detected, by RT-qPCR, the presence of cytokine and chemokine mRNA in O4+ OPCs in control conditions (Figure 6A). These mRNA levels were greatest at P3 and significantly decreased in a stage-dependent manner, between P3 and P10 (Figure 6A). In addition, we verified that our control condition (i.p. PBS) did not constitute a stress, *per se*, that would induce the expression of cytokine and chemokine genes. Comparing MACS-isolated O4+ OPCs from naïve (untreated) and PBS-treated pups at P5, we observed similar levels of cytokine and chemokine gene expression, in RT-qPCR experiments (Figure S7B). This shows that PBS injection is not responsible for constitutive cytokine and chemokine mRNA levels at P5. In contrast, OPCs isolated from IL1B-treated mice exhibited elevated levels of cytokine and chemokine mRNAs compared to naïve or PBS OPC samples, as expected (Figure S7B). Moreover, we confirmed that the constitutive expression of cytokine and chemokine genes also occurred in normal conditions in the murine oligodendroglial cell line, Oli-neu (Figure 6B,C), therefore indicating that the detection of such mRNAs is neither due to contamination of O4+ OPCs by MG, nor to their isolation mode, nor to the intraperitoneal injection *per se*. We also showed that this constitutive expression decreased during Oli-neu differentiation – using two protocols – as it was observed during the O4+ OPCs maturation trajectory (Figure 6C and Figure S1E). These data demonstrated that O4+ OPCs intrinsically transcribe cytokine and chemokine genes at an early OPC stage (P3), and that the expression of these genes is gradually downregulated during their maturation process between P3 and P10, in a physiological and developmental manner. This constitutive transcription is in line with the small extent of chromatin accessibility changes of these genes that we observe at P5 (Figure 2F).

**Figure 6.**
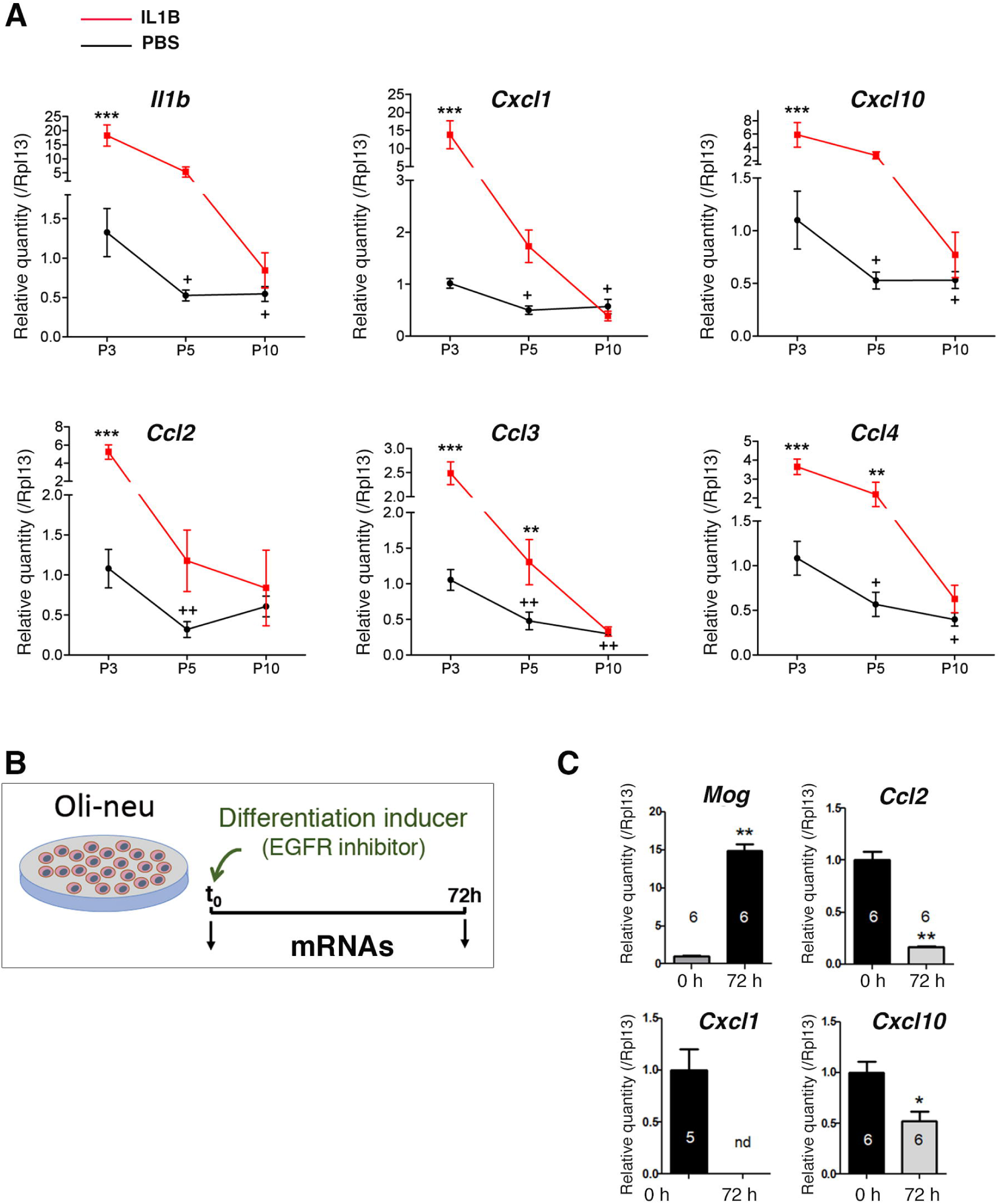
Constitutive expression of genes encoding cytokines and chemokines at early stages of normal (unstressed) O4+ OPC maturation. (see Figures S1E and S7). (**A**) RT-qPCR analyses in P3, P5, and P10 O4+ OPCs. N= 5 to 6 independent experiments. Two-way ANOVA followed by Bonferroni - Post Hoc Test was performed: * and +, p < 0.05; ** and, ++ p < 0.01; ***, p < 0.001. *, **, or ***, correspond to comparison between PBS and IL1 conditions for a given postnatal stage; + (or ++), correspond to comparison between P5 (or P10) to P3. (see Figure S7). (**B** and **C**) ***The oligodendroglial cell line Oli-neu reproduces the constitutive expression of cytokine and chemokine gene at the immature state and its downregulation upon differentiation*** (see Figure S1E) (**B**) Differentiation protocol of the Oli-neu cell line triggered by PD174265, a potent, cell-permeable inhibitor of the tyrosine kinase activity of the epidermal growth factor receptor (EGFR) (**C**) RT-qPCR analyses in the Oli-neu cell line before or after 72 hours of differentiation. This *in vitro* paradigm shows the same pattern as O4+ OPCs, in terms of inflammatory mediator gene expression at the immature (undifferentiated) state and decrease when maturating to myelin-producing oligodendrocytes. Numbers of independent experiments are indicated on the graph bars. *, p < 0.05; **, p < 0.01. nd: not detected.

To further confirm our assumption that, in OPCs, at P5, the chromatin of neuroinflammation-induced genes exhibited “neuroinflammatory-like” signatures, prior to IL1B-treatment, we formulated and performed an additional analysis, taking advantage of an existing public ATAC-seq dataset from a similar control versus treatment study, which used IL1B stimulus on human adult aortic endothelial cells (HAECs) isolated from aortic trimmings of donor hearts (Hogan et al., 2017; “HAEC dataset”; NCBI Gene Expression Omnibus; accession no: GSE89970). Through a cross-species comparison, our aim was to explore global chromatin landscape similarities (or lack thereof) between both control and IL1B-treated, in adult (HAEC) versus neonates (O4+ OPC) samples. First, both datasets were limited to chromatin regions annotated with matching 1-to-1 gene orthologs and located within ±2 kb of a TSS as described in Material and Methods. In total, we were able to match 7,739 peaks between the two datasets, including 100 regions, which were found to be different between the control and IL1B conditions, in the HAEC dataset. Subsequent cross-comparison of these regions reported no significant difference between both O4+ OPC samples and the HAEC IL1B-treated sample. In contrast, both O4+ OPC samples showed significant differences (p-value < 10^−15^) to the HAEC control sample (Figure S7C,D; Table S10). These findings reveal a characteristic “neuroinflammatory-like” pattern in the chromatin profiles of OPCs isolated from both inflammation-exposed and control mice, which is indicative of an inflammation-like signature that is already present at basal level in developing OPCs. The results thereby reinforce our findings about cytokine and chemokine gene expression in unstressed and stressed O4+ OPCs.

### Correlative upregulation of genes of the immune/inflammatory pathway and downregulation of the myelination program in an oligodendroglial cell line upon inflammatory stimulus

Our results suggest that 1) there is a specific and restricted time-window for the endogenous expression of the cytokine and chemokine genes along the OPC maturation trajectory; 2) that, in response to *in vivo* IL1B-induced neuroinflammation, which is known to result in a blockade of OPC maturation (Favrais et al., 2011), the overexpression of the genes of the immune/inflammatory pathway strikingly correlates with the reduction of the expression of the myelination programme *in vivo* (Figure 6A; Figure S7A). This is recapitulated in an oligodendroglial cell line *in vitro* (Figure 6B,C and Figure S1E). To further explore the link between these two processes, we investigated the response of the Oli-neu cell line to an inflammatory stimulus during its differentiation time-course. We showed that *ex vivo* treatment of the Oli-neu cell line by TFN-alpha compromised the expression of genes of the myelination programme - as expected -, which is a hallmark of differentiation blockade (illustrated by *Cnp, Mbp* gene expression; Figure 7). In addition, it also induced concomitant strong and abnormal overexpression of genes of the immune/inflammatory pathway, thereby counteracting the physiological decrease of cytokine/chemokine expression normally observed during differentiation (Figure 7; here, illustrated for *Ccl2 and Cxcl10*). These data indicate that exacerbating the immune/inflammatory phenotype would participate to differentiation blockade.

**Figure 7.**
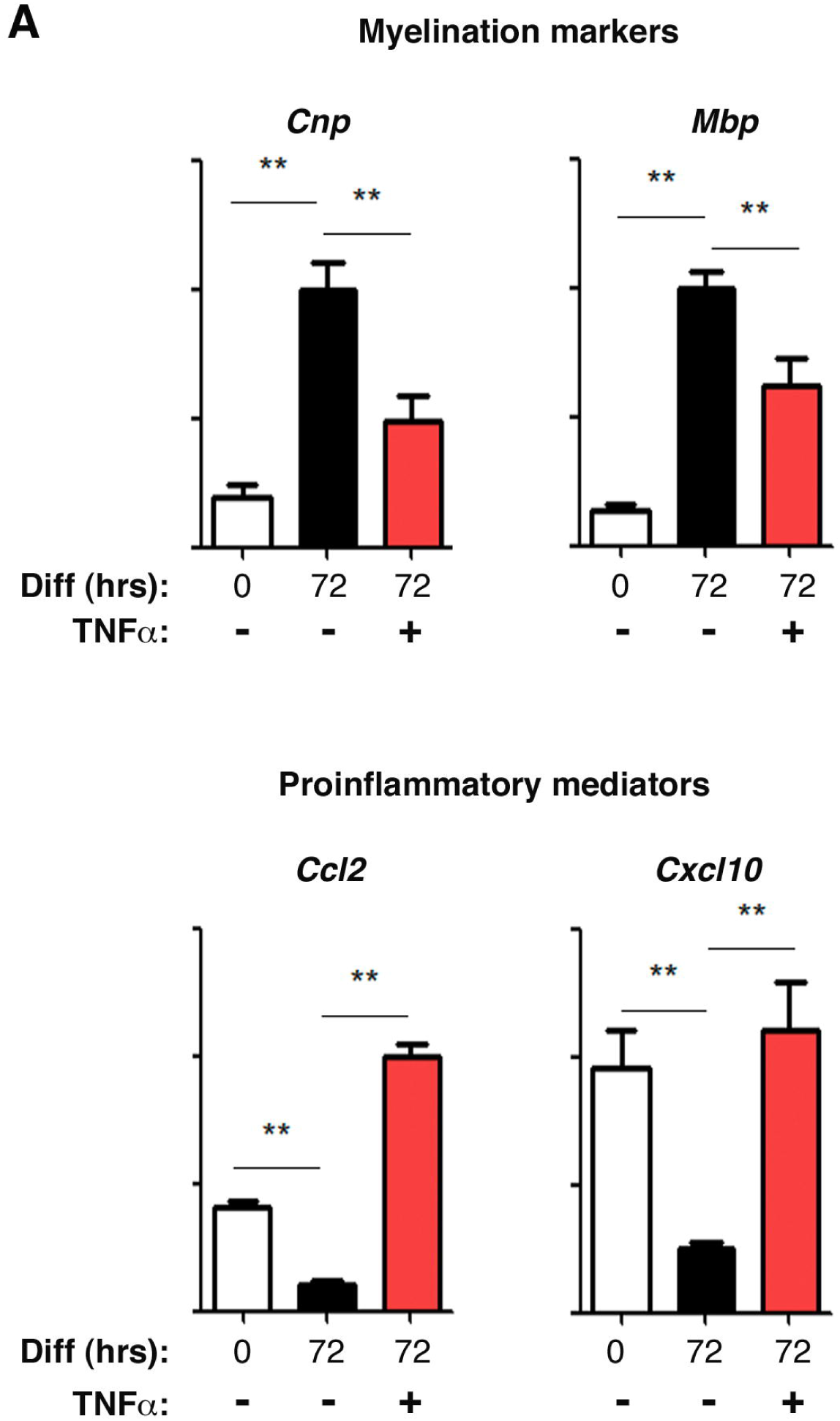
Correlative upregulation of genes of the immune/inflammatory pathway and downregulation of the myelination program in an oligodendroglial cell line upon inflammatory stimulus. *Ex vivo* treatment of the Oli-neu cell line by TFN-alpha reproduces the disturbances in gene expression that we observe in isolated O4+ OPCs upon *in vivo* neuroinflammation. Illustrated here for the *Cnp, Mbp*, genes, and *Ccl2 and Cxcl10*, respectively. RT-qPCR analyses (n = 6 independent experiments). Oli-neu cells were differentiated by 1 µM of PD174265 as in Figure 6C,D (Gobert et al., 2009; see materials and methods), during 72h, in absence (T72 co) or in presence of 10 ng/ml TNFα (T72 +TNFa). T_0_: undifferentiated cells. Non parametric t-tests; ** p< 0.01.

## DISCUSSION

The main goal of our study is to decipher molecular mechanisms that are triggered by neuroinflammation in a context leading to OPC maturation arrest and white matter injury. Most of the studies on prenatal stress has focused on the disturbances of epigenetic marks and has not really addressed the robustness versus vulnerability of the epigenome, towards these insults (Schang et al., 2018). Moreover, they were performed on the whole brain or dissected regions. There are therefore two main interests in our study.

The first major interest of our study is to use a purified population of OPCs, based on the O4 marker to investigate the respective epigenomic and transcriptomic contributions to the disturbances induced by neuroinflammation. To do so, we carefully assessed the purity of our isolated populations by different means and at each step of our study, as summarized in Figure S1. Globally, we showed that specific markers of MG (and astrocytes; Figure 4) are unexpressed in the isolated O4+ OPC population, based on transcriptomic and RT-qPCR data. Moreover, the chromatin conformation of the corresponding MG-specific genes is inaccessible, based on ATAC-Seq data, in contrast to that of chromatin regions of genes characteristic of OPCs. These data indicate that our O4+ population is highly enriched in OPCs. Moreover, we could recapitulate in an oligodendroglial cell line the two major aspects of normal and perturbed OPC maturation that we unraveled in O4+ OPCs: 1) the constitutive expression of cytokine and chemokine genes in unstressed conditions and its physiological downregulation during differentiation; 2) the upregulation of these genes in response to inflammatory insult. In addition, these genes encoding cytokines and chemokines are in a closed chromatin conformation in MG, whereas they are in an open conformation in both unstressed and stressed OPCs. Therefore, the assessment of the purity and functionality of this isolated O4+ OPC population is built on a corpus of arguments that altogether lead to the conclusion that O4+ OPCs are able to constitutively synthesize molecules belonging to the immune/inflammatory pathways.

The second interest of our study is to show that, surprisingly, the epigenome of OPCs is globally preserved in response to neuroinflammation, since only a very restricted number of regions undergo opening or closing of the chromatin. Rather, our results indicate that neuroinflammation has a major impact at the transcriptomic level, by disturbing the transcription of genes that are already active in OPCs at the time of exposure, under unstressed conditions. Indeed, one route of entry for neuroinflammation is the remarkable capacities of premyelinating O4+ OPCs to produce immunomodulators. More precisely, the developmental control of their synthesis – namely, expression at P3 and downregulation at P5 and P10 – is highjacked by neuroinflammation, which provokes abnormal overexpression of these genes at a stage at which they should be downregulated. As such, this process occurs with limited effects on chromatin accessibility. In other words, rather than remodeling the epigenetic landscape, the response to neuroinflammation takes advantage of and place in a “primed” epigenetic landscape. Such a mechanism is reminiscent of what happens in response to cellular stress, involving the heat shock pathway driven by the Heat Shock transcription Factors, which is “pre-wired” by the chromatin architectural characteristics (HSFs; Vihervaara et al., 2017; Vihervaara et al., 2018). Interestingly, a similar situation has been observed during learning and memory processes, where increase in the levels of histone posttranslational modifications seem restricted to genes that are already active in the naive state, while, conversely, global histone modifications are vastly uncoupled from changes in gene expression (Halder et al., 2016).

In line with the expression of genes of the inflammatory pathway in OPCs both under unstressed and stressed conditions, we found that, globally, characteristic binding sites for TFs involved in the immune/inflammatory pathway are occupied before and after exposure to neuroinflammation. However, our finding of existing paired binding motifs for these TFs, in ATAC-Seq peaks, suggest that these factors can act in a combinatorial mode and that changes in their combination (and/or activity, as observed in other contexts; Mancino et al., 2015) might account for the increase in the transcription of the corresponding genes that we detect upon neuroinflammation exposure.

Our data also nurture an underlying and emerging concept: molecules, which have been historically identified and studied as key mediators of stress responses and guardians of cell or organism homeostasis, are also pivotal in physiological conditions for normal development (Cardona et al., 2008). Emblematic examples are represented by TF families, like NFkB (Espín-Palazón and Traver, 2016), HSFs and their target genes encoding the heat shock proteins (HSPs; Abane and Mezger, 2010; Chang et al., 2006; Gomez-Pastor et al., 2018), and by critical players of the unfolded stress response (UPR; Laguesse et al., 2015). Whether these two apparently distinct functions have emerged concomitantly in evolution or not is unclear, but for technical and practical reasons, the roles of these molecules in normal development have been understudied. The stage-dependent production of inflammatory players by prenatal OPCs during the time-course of their maturation, as well as their potential role in brain formation, is therefore less unexpected than it appears. Accordingly, it has been extensively reported in the literature that OPCs and mature oligodendrocytes of the adult brain can express key players of immune and inflammatory pathways in pathological conditions. This includes studies of patients affected by multiple sclerosis (MS) or in *in vivo* models of experimental autoimmune encephalomyelitis (EAE; reviewed in Zeis et al., 2016). Here, we show that P5 O4+ OPCs express genes and proteins associated with inflammatory pathways in normal conditions. As such, OPCs during normal development thus display properties similar to that ascribed to adult OPCs and mature oligodendrocytes, which can shape the inflammatory environment, or - in the perspective brought by the new concept mentioned above - could perform a trophic role on their environment at defined time-windows. Our results are also in line with previous results showing that OPCs, derived *in vitro* from neurospheres, can activate cytokine genes in an EAE model (Cahoy et al., 2008). In a more original manner, we also unravel the physiological, constitutive expression of cytokine and chemokine genes in normal O4+ OPCs at an early postnatal stage. In line with our findings, Zeis et al. (2016) revisited published microarray data sets (Cahoy et al., 2008) and pointed out the expression of genes belonging to the GO:term “immune system process”, in PDGFRa+ unstressed OPCs, which reinforces our data.

Among the genes belonging to the immune system and inflammatory pathways and undergoing dysregulated expression upon exposure to neuroinflammation, only a limited number exhibit significant chromatin remodeling. Hypoxia-Inducible Factor 3, *Hif3a*, is one of them and was shown to be regulated, in an oxygen-independent manner, in two distinct models of inflammation, in non-neural cells (Kumar et al., 2015; Cuomo et al., 2018). Interestingly, in parallel of our data, Cuomo et al. (2018) established that proinflammatory cytokines are responsible for the activation of *Hif3a* gene, through epigenetic changes and the involvement of NFκB.

One question is the functional impact of the opening or closing of the chromatin in response to neuroinflammation, in regions that we have identified by ATAC-Seq, since most of them do not correlate with major transcriptional changes. This raises three interpretations. Firstly, the transcriptomic impacts of these epigenomic modifications might be “buffered”, thanks to the binding of different sets of TFs, which would deserve further investigations. Secondly, TFs, which are multifaceted drivers, remodel the chromatin state and genome topology, often before changes in gene expression can be observed, as was demonstrated in studies on the molecular basis of cell fate (Stadhouders et al., 2018). These modifications of the chromatin landscape could thus constitute an Achilles’ heel for transcriptomic disturbances, that would occur either at later maturation stages, at temporal distance from the insult, or upon a second hit of neuroinflammation. The occurrence of additional inflammatory insults is relevant because, besides exposure to prenatal inflammatory insults - which is mimicked by our model -, preterm babies also face a heavy burden in terms of postnatal inflammatory insults, representing additional neuroinflammatory hits (Barnett et al., 2018). Thirdly, one other exciting possibility is that, in the genome, these TFs involved in immune and inflammatory pathways might also work at long-distance from their dysregulated target genes, which would involve remodeling at multiple architectural levels (chromatin looping, sub-TADs (topologically associated domains) connectivity etc.).

Finally, besides these considerations, the entry route for neuroinflammation is represented by the constitutive and stage-dependent synthesis of cytokines and chemokines by premyelinating OPCs in normal conditions, which empowers neuroinflammation to impact OPC maturation. This entry route can thus be envisioned as recapitulating and intermingling both injurious and developmental aspects, as already expected from the field (Volpe, 2011; Deverman and Patterson, 2009; Mancino et al., 2015). Indeed, by counteracting the tightly regulated physiological expression of cytokines and chemokines by O4+ OPCs at P3, that is programmed to gradually decrease in a developmental, stage-dependent manner (here shown between P3 and P10), the neuroinflammatory insult might compromise the premyelinating OPC cell fate. Although the causality of the abnormal upregulation of immune/inflammatory modulators by OPCs on their maturation is difficult to prove in our system, it is strongly supported by an array of arguments: First, we can reproduce the correlation between the upregulation of cytokine and chemokine gene expression and the downregulation of the myelin markers in an oligodendroglial cell line, upon proinflammatory exposure to TNF-alpha. Second, Moyon et al. pointed out the role of IL1B and CCL2 production by premyelinating OPCs in modulating their motility capacities and eventually differentiation (Moyon et al., 2015). Third, CXCR2 counteracts adult OPC differentiation and myelination potential, in a model of multiple sclerosis, by interfering with the PI3K/AKT/mTOR pathway, thereby strongly reinforcing the possibility of a role of cytokines and chemokines in OPC maturation during development (Wang et al., 2020). One possible pathway for the counteracting effects of cytokines and chemokines on OPC differentiation is that their production could regulate the recruitment by OPCs of other cells that are known to influence OPC maturation (like microglia; (Jana and Pahan, 2005; Balabanov et al., 2007).

In conclusion, in the context of a chronic perinatal systemic inflammation, the epigenome seems globally preserved in premyelinating OPCs, in terms of chromatin accessibility, suggesting that the contribution to OPC blockade is mostly driven by transcriptomic disturbances. These transcriptomic perturbations concern transcriptional programs that are already active at the time of exposure, and, most prominently, those involved in the immune system and inflammatory pathways. Our results have important therapeutic consequences: because of the striking intertwining between the injurious and developmental facets of these inflammatory mediators, we should reconsider that global targeting of this pathway might constitute a therapeutic option. One conclusion emerging from our study is that the TFs involved in response to neuroinflammation in OPCs seem to be already at play in normal OPCs to control the developmental transcription of these genes. In addition, some of them work in combination as suggested by their pairing profiles. Our work thus paves the way of future studies that would allow the design of a therapeutic strategy, based on subtle manipulation of the activity of TFs, using appropriate cocktails of low-dose modulators.

## MATERIALS AND METHODS

### Animal Model

Experimental protocols were approved by the institutional review committee (under the following reference by the French Ministère de l’Enseignement Supérieur et de la Recherche (#2016040414515579) and met the guidelines for the United States Public Health Service’s Policy on Humane Care and Use of Laboratory Animals (NIH, Bethesda, MD, USA). Sex was determined at birth, and confirmed by abdominal examination at sacrifice. This animal model is similar to the human in that males are more affected and, as such, only male OF1 pups were used, as female OPCs maturation is not altered (Hagberg et al., 2015). IL1B injections were performed as described (Favrais et al., 2011; Krishnan et al., 2017). Five µL volume of phosphate-buffered saline (PBS) containing 10µG/kG/injection of recombinant mouse IL1B (R&D Systems, Minneapolis, MN) or of PBS alone (control) was injected intraperitoneally (i.p.) twice a day on days P1 to P4 and once a day, on day P5 (see Figure 1). Pups were sacrificed four hours after the morning injection of IL1B at P3 or P5, and at a similar time at P9, or P10. ATAC-Seq data were produced from 3 independent biological replicates for each condition (PBS or IL1B; Figure 1). Microarray data were produced from 6 independent biological replicates for each condition (PBS or IL1B; Figure 1), using the same animals that were also analysed for CD11B+ microarrays (Krishnan et al., 2017).

### O4+ and CD11B+ microglial magnetic activated cell-sorting in mouse

O4+ cells were isolated at P3, P5, P9, or P10 by Magnetic Activated Cell Sorting (MACS, Miltenyi Biotec, Bergisch Gladbach, Germany), according to the manufacturer’s protocol and as previously described (Schang et al., 2014). Briefly, brains were collected without cerebellum and olfactory bulbs, pooled (3 brains per sample) and dissociated using the Neural Tissue Dissociation Kit containing papain. O4 + cells were then enriched by MACS, using the anti-O4 MicroBeads. For microarray and RT-qPCR analysis, the eluted isolated cells were centrifuged for 5 min at 600g and conserved at −80°C. CD11B+ microglial cells were isolated as described (Krishnan et al., 2017). The unlabeled fraction mainly contained astrocytes (see Figure S1A). For the ATAC-seq experiment, 50,000 cells were immediately lysed and their nuclei submitted to Tn5 activity. The purity of the eluted O4-positive fraction was verified using qRT-PCR for Myelin Basic Protein (*Mbp*), ionizing calcium binding adapter protein (*Iba1*), glial fibrillary acid protein (*Gfap*) and neuronal nuclear antigen mRNAs (*NeuN*; Figure S1A). Comparable numbers of O4+ OPCs from control (PBS) and treated (IL1B) samples were collected (1.12 × 10^6^+/- 0.12 × 10^6^ cells per sample).

### OPC culture and differentiation

OPCs were prepared from newborn OF1 mice as described (McCarthy and de Vellis, 1980; Pansiot et al., 2016). In brief, forebrain cortices were removed from postnatal day 0–2 mouse pups and freed from meninges. Minced tissues were enzymatically digested with 0.125% trypsin (Sigma) and 0,0025% DNase I (Sigma) for 15 min at 37°C and then mechanically dissociated. Cells were filtered through a 100-μm-pore-size cell strainer (BD), centrifuged 10 min at 1800 rpm, resuspended in minimum essential Eagle’s medium (Sigma) supplemented with 10% FBS (Gibco), 1% Glutamax (Gibco), 1% penicillin-streptomycin (P/S) solution (Sigma), and 0.5% glucose and plated in T75 flasks at a density of 2 × 10^5^/cm^2^. Mixed glial cell cultures were grown until confluence for 9-11 days (medium was replaced every 48-72h) and shaken for 1.5 h at 260 rpm to detach microglia. These detached microglia were then collected and removed together with the media. Remaining cells were shaken for additional 18h to detach the OPCs from the astrocyte base layer, and were simultaneously treated with 100 µg/ml liposomal clodrosome suspension (Clodrosome®, Encapsula Nanosciences, Brentwood, USA) which selectively eliminates any residual microglia. The detached OPC cell suspension was filtered through a 20-µM-pore-size filter (Millipore) and incubated in an untreated Petri dish for 10 min at 37°C to allow attachment of any remaining microglia. Purified OPCs were then seeded onto poly-D-lysine-coated 12-multiwell plates at a density of 3 × 10^4^/cm^2^ in OPC proliferation medium composed of Neurobasal medium (Gibco), 2% B21 (Miltenyi biotec), 1% P/S (Sigma) and 1% Glutamax (Gibco), supplemented with growth factors consisting in 10nG/mL FGFα (Sigma) and 10nG/mL PDGFα (Sigma). After 72h, OPC differentiation was initiated by growth factor withdrawal and addition of 40 nG/mL of T3 (Sigma). At the same time, OPCs were treated with 50nG/mL IL1B (R&D Systems, Minneapolis, MN) or PBS for 4h, treatment was removed, new media provided and cells were grown in differentiation medium until 72h (Figure 4C).

### Oli-neu cell line culture and differentiation

The immortalized murine OPC cell line, Oli-neu, was kindly provided by Dr Sheila Harroch (Pasteur Institute, Paris, France). Oli-neu was established from OPC-enriched murine primary cultures from E16 brains transformed with a provirus containing the oncogene *T-Neu* (Jung et al., 1995). Various differentiation protocols have been established, among which treatment with PD174265, a selective inhibitor of the activity of Epidermal Growth Factor receptor (ErbB) tyrosine kinase, has been shown to induce MBP expression (Gobert et al., 2009). These cells were cultured in Dulbecco’s modified Eagle’s minimum essential medium (DMEM) containing Glutamax 1X and high glucose (4.5 G/L; Gibco 31966), supplemented with 1 mG/mL insulin (Sigma), N2 supplement (Gibco), 100 μG/mL T4 and T3 (Sigma), 1% horse serum (Gibco), and 1% P/S (Sigma). At confluence, the cells were mechanically detached and seeded in 12-multiwell plates at a density of 3 × 10^4^ cells/cm^2^. After 24h, differentiation was induced by addition of 1µM PD174265 (ChemCruz) diluted in DMSO at 1 mM. Medium was replaced after 48h and differentiation was stopped after 72h (Figure 6B,C, Figure S1E).

### RT-qPCR analysis and Luminex assay

Preparation of samples for quantitative reverse-transcriptase polymerase-chain reaction (qRT-PCR), primer design PCR protocol and luminex assay were similar to that previously described (Chhor et al., 2013). Primer sequences are given in Table S11. *Gapdh* (glyceraldehyde-3-phosphate dehydrogenase gene) and *Rpl13* (Ribosomal Protein L13) were chosen to standardize the quantitative experiments based on reference gene suitability testing.

### ATAC-Seq analysis in O4+ OPCs

ATAC-seq protocol was performed as described (Buenrostro et al., 2013) with slight modifications. In brief, cells were immediately lysed after cell sorting and a total of 50,000 nuclei were subjected to Tn5-mediated transposition for 30 min, resulting in ‘tagmented’ DNA fragments. Tagmented DNA was purified on MinElute colums (Qiagen) and amplified/tagged in two steps using NEBnext High-Fidelity 2x PCR master mix (New England Biolabs). Amplified DNA was purified twice with 1.8 volumes of NucleoMag NGS Clean-up and Size Select beads (Macherey Nagel). DNA was quantified using the Qubit dsDNA HS Assay Kit and the quality of each library determined on Agilent 2100 Bioanalyzer DNA High Sensitivity ChIPs. Libraries demonstrating appropriate nucleosomal profiles were multiplexed and subjected to Illumina NextSeq500 sequencing (IGenSeq Platform, ICM, Paris, France). Fastq files are available in Dataset S1. The main steps of sequence analyses are summarized in Figure S2B and detailed on the jupyter notebook available in supplemental information. After quality controls (Fastqc and Trimmomatic 0.33), reads were aligned on the mm10 genome with Bowtie 2 (Galaxy tool version 2.3.4.1 (Afgan et al., 2016); default parameters; Table S1; Figure S2B). Peak calling was performed with MACS2.2.0; default parameters; q<0.05) separately for the two conditions, using a pooled (n=3) bam file of control samples and a pooled (n=3) bam file of IL1B samples. The two resulting bed files were merged and, after removing the mm10 blacklist (http://mitra.stanford.edu/kundaje/akundaje/release/blacklists/mm10-mouse/mm10.blacklist.bed.gz), 213,246 DNA regions (peaks) significantly detected in at least one condition were delimitated (Table S2). The number of reads was determined in each peak for each sample using Bedtools coverage (version 2.19.1) and normalized to the library sizes. Principal component analysis was performed on log transformed read count values of the top 500 most variable peaks, using the prcomp function in R. Differential peak detection between the three PBS and the three IL1B samples was performed with the Bioconductor software package EdgeR (3.22.3; Robinson et al., 2010), using R studio (0.98.1103; http://www.rstudio.com). Statistical comparison was performed using the exact test function followed by False Discovery Rate (FDR) determination by the Benjamini-Hochberg method. Raw data are available under SRA BIOPROJECT accession # PRJNA540409.

### Linking of HAEC and OPC ATAC-Seq datasets

We used a public ATAC-Seq dataset of human aortic endothelial cells (HAECs; Hogan et al., 2017; NCBI Gene Expression Omnibus; accession no: GSE89970) and processed the raw reads (using the hg19 reference genome) to obtain a set of peaks. Both sets of peaks (control and IL1B-treated samples) were annotated using HOMER’s annotatePeaks function. Next, HAEC peaks were matched to mouse OPC peaks through gene annotations, by taking only those peaks annotated with matching orthologous genes (only 1-to-1 orthology was considered). Matching was further restricted to promoter regions (peaks with a relative maximum distance of 2kb from the TSS). In order to ensure that peaks were true matches, this set was further restricted to a relative distance of 500 bp from each other in relation to the TSS. Using this approach, a total of 7,739 peaks were matched between the HAEC and OPC datasets, including 100 peaks identified as differential in the HAEC dataset using DESeq2. Next, the number of reads mapped to matched peaks were obtained by counting the number of reads at the summit ± 50bp using the featureCounts package of the Subread software (v1.6.0) and the counts were normalized against the total number of reads present in all matched peaks and converted into reads per million. Normalized read number distributions of the two datasets were compared using the one-sample Wilcoxon rank test with continuity correction.

### Microarrays of mouse O4+ OPC gene expression and data preprocessing

Microarray analysis was performed on six control and six IL1B samples (O4+ cells isolated at P5 after *in vivo* PBS or IL1B treatment) using Agilent Whole Mouse Genome Oligo Microarrays 8×60K (Agilent). Raw data are available in Dataset S2. All the steps, from RNA extraction to statistical analysis, were performed by Miltenyi Biotec, as previously described (McCarthy and de Vellis, 1980). In brief, intensity data were subjected to quantile normalization, unpaired t-tests (equal variance) were conducted to compare intensities between the two groups for each probe and p-values were adjusted through FDR determination by the Benjamini-Hochberg method. Fold changes correspond to the median ratios (median[IL1B]/median[PBS]). When FC<1, the FC was expressed as a negative value using the formula FC(neg)=-1/FC. For example, if FC=0.5, the indicated FC is −2. Probes with FDR <0.05 were considered significant. An additional fold change (FC) threshold was chosen at +/- 1.5 (corresponding to FC>1.5 and <0.666).

### Heat map representation

Heat maps were created using Morpheus (https://software.broadinstitute.org/morpheus). The Log2 median-centered data were visualized using a fixed (nonrelative) color pattern. The color scales are indicated on each heatmap. Rows and columns were submitted to hierarchical clustering with the following criteria: metric = one minus Pearson correlation, linkage method = average.

### GO-term enrichment analysis

GO-term Biological Pathway enrichment was done using David 6.8 (Huang et al., 2009a; Huang et al.,

2009b).

### TFBS motif enrichment analysis, and TF footprint analysis

The 213,246 significant peaks detected by MACS2 in at least one condition (PBS or IL1B) were annotated with the *HOMER annotatePeaks* function. The list was restricted to the peaks located between −8000 and +8000 bp from the closest TSS (“TSS-All” list). Among this list, peaks were selected, which were annotated with a gene name and for which the gene expression was modulated in the microarray analysis (FDR<0.05 and FC>1.5 or <-1.5). The full list of peaks and lists restricted to up or down-regulated genes were submitted to motif enrichment analysis using *HOMER FindMotifsGenome* with the options “-size given” and “-mask”. The “TSS-All” list was used as background. Six motifs corresponding to the top 5 motifs enriched in the full list of peaks (ISRE, IRF1, IRF2, Nfkb-p65 and PGR, Figure S5B) and the 15th motif enriched in the list restricted to up-regulated genes (PU1:IRF8, Figure S5B) were localized in the full list of peaks (UP+DOWN) with *HOMER FindMotifs*. For each of these motifs, the average profile of Tn5 activity was visualized using *pyDNAse* dnase_average_profile.py; Piper et al., 2013). This profiling was performed using a pooled bam file of PBS samples, or a pooled bam file of IL1B samples separately, and a pooled bam file of the two conditions (“both”) together.

### Testing for enrichment of paired motifs

2319 ATAC-Seq peaks corresponding to 1266 upregulated genes (886 different gene names) and 946 ATAC-seq peaks corresponding to 454 downregulated (336 different gene names) were tested for significantly enriched pairs of TFBS relative to a universe containing all the peaks located +/- 8 kb around the closest TSS. For each individual motif from the homer database, all peaks in the universe were ranked by motif occupancy using a binomial score. Then for every possible pair of motifs, peaks containing both motifs were identified using the overlap between top 5000 ranked peaks for each of the individual motifs. A hypergeometric test was used to calculate the enrichment score (p-value) for the overlap between each test set and the peaks containing both motifs. The resulting p-values were corrected using the Benjamini-Hochberg correction.

### Statistical analysis

All *in vivo* and *in vitro* experiments were performed using an alternating treatment allocation. All analyses were performed by an experimenter blinded to the treatment groups. The results of qRT-PCR and Luminex analyses are expressed as mean +/- SEM of at least four independent experiments; the number of analysed samples is indicated in the figure legends or on the graphs. Statistical analysis was done using the non-parametric Mann-Whitney t-test with Graphpad 5.0 software (San Diego, CA, USA) or two-way ANOVA followed by Bonferroni - Post Hoc Test as indicated in each figure legend. Significance is shown on the graphs (*, p < 0.05; **, p < 0.01; ***, p < 0.001). Specific statistical analyses for ATAC-seq and microarray analyses are detailed in the dedicated sections of Material and Methods. The significance of intersection between the two datasets was evaluated by hypergeometric test (Phyper function) in R studio.

## LEGENDS OF SUPPLEMENTAL FIGURES

**Figure S1. Assessment of the purity of the isolated O4+ OPC population** (related to Figures 1-4,6).

(**A**) ***Quality assessment of the O4+ OPC cell purification process*** (relative to Figure 1) RT-qPCR experiments on the O4+ OPC, CD11B+ microglia (MG; Krishnan et al., 2017; n=3 PBS samples and n=3 IL1B samples) and unlabeled cell populations, showing that, in contrast to MG (CD11B+) and astrocytes (unlabeled), the population of O4+ OPCs used in this study express the *Mbp* gene, whereas it exhibits very low mRNA levels of the microglia marker CD11 (encoded by the *Itgam* gene), the astrocyte marker *Gfap*, or of the neuronal marker *NeuN*. Note that *NeuN* is very lowly expressed, even in the unlabeled population, which mainly contains astrocytes, because neurons poorly survive our MACS protocol.

**(B) *Distinct perturbations in gene expression profiles in isolated O4+ OPCs and CD11B+ MG, at P5 upon neuroinflammation***. Heat maps of the overall comparison of neuroinflammation-induced transcriptomic changes. (related to Figure 4). OP4+ OPCs exhibit profiles of transcriptomic modifications globally very different from that of CD11B+ microglial cells (data by Krishnan et al., 2017). Notably, these cell populations were <u>isolated from the same animals at P5</u>. Examples of genes encoding cytokines and chemokines are pointed out by rectangles. Most of them are not upregulated in CD11B+ microglial cells at P5, in contrast to what happens in O4+ OPCs. Log2 Fold change ([-2.0; +2.0]; IL1B/PBS)

(**C**,**D**) ***Distinct profiles of chromatin accessibility in genes specific for OPCs*** (our ATAC-seq data, green peaks; Figure 2 and Figure S2,3) ***and MG*** (ATAC-Seq data mining from (Matcovitch-Natan et al., 2016; red peaks).

(**E**) ***The oligodendroglial cell line Oli-neu reproduces the constitutive expression of cytokine and chemokine gene at the immature state and its downregulation upon differentiation observed along the maturation time-course***. (relative to Figure 6B,C). RT-qPCR analyses in the Oli-neu cell line before or after 72 hours of differentiation triggered by exposure (+) to conditioned medium (CM) from primary neuron culture.

**Figure S2. ATAC-Seq data quality control**. (related to Figure 2)

(**A**) ***Quality-control validation of OPC maturation arrest in the model***. RT-qPCR analysis of the expression of myelination and progenitor markers in OPCs. <u>Myelin markers</u>: *Mbp*, Myelin binding protein; *Mog*, Myelin oligodendrocyte glycoprotein; *Mag*, Myelin-associated glycoprotein. *Plp1*, proteolipid protein 1, a transmembrane, predominant component of myelin; *Cnp*, 2’,3’-Cyclic Nucleotide 3’ Phosphodiesterase, abundant protein in myelin in the central nervous system. Progenitor (OPC) markers: *Id2*, Inhibitor of differentiation 2; *Pdgfra*, Platelet Derived Growth Factor Receptor Alpha. Number of independent experiments: n = 7 for *Cnp, Mag*, and *Pdgfra*; and n = 15 for *Mbp, Mog*, and *Id2*. ns: not statistically significant; *, p < 0.05; ***, p < 0.001.

(**B**) ***Schematic representation of the bioinformatics and statistics workflow used for the analysis of ATAC-Seq data***

(**C**) ***Fragment-length distributions in ATAC-Seq samples***. Insert size distribution shows visible large periodicity of the nucleosome-free, mononucleosomal and dinucleosomal fragments, as well as the expected ∼10.4bp periodicity, resulting from steric hindrance of the helical twist of the DNA on the nucleosome surface.

**Figure S3. The epigenome of OPCs is globally preserved after IL1B treatment (ATAC-Seq analysis)** (related to Figure 2)

(**A**) Multidimensional Scaling (MDS) plot of distances between the 3 PBS samples and the 3 IL1B samples, using EdgeR.

(**B**) Scatter plots representing the dispersion (logfold change) as the number of tn5 cuts per million (logCPM), for each individual analyzed peak (across 3 PBS samples and 3 IL1B samples). In red, peaks showing differential chromatin accessibility with FDR < 0.05. The upper left plot is <u>identical to Figure 2B</u> and is included here for comparison. Upper right, lower left, and lower right: scatter plots for analyses with permuted sample labels, created by swapping of one of the PBS samples with one other IL1B samples or, vice-versa. Mixing PBS and IL1B samples led to very reduced numbers of differential peaks, confirming that differential peaks found with correct sample labels are not statistical artefacts

(**C**) Distribution of the total number of peaks and of the 524 differential open peaks (right-hand side) in various genomic regions.

(**D**) GO analysis, using DAVID6.8 (Huang et al., 2009a; Huang et al., 2009b), using genes with TSS of genes that are annotated with one or more differentially accessible chromatin regions (total of 524 ATAC-Seq peaks).

**Figure S4. The induction of cytokine and chemokine mRNA levels, upon IL1B exposure, is reproduced in MACS-isolated *ex vivo* cultured OPCs during differentiation**. (related to Figure 3 and Figure 4C,D)

(**A**) ***Quality control of the samples used for microarray analysis***. Relative expression, by microarray analysis, of several myelination markers and Inhibitor of Differentiation *Id2* in O4+ OPCs at P5 (6 PBS samples and 6 IL1B samples)

(**B**) ***GO analysis corresponding to the TOP-10 genes, which are downregulated upon IL1B treatment, using DAVID6***.***8***.

(**C**) ***Cultured O4+ OPCs express cytokine and chemokine genes***. RT-qPCR experiments on OPCs collected after 72h of differentiation (see experimental design on Figure 4C). Number of independent experiments: n = 6 per condition (PBS or IL1B). *, p < 0.05; **, p < 0.01; ***, p < 0.01.

**Figure S5. TFBS motifs identified in significant ATAC-seq peaks** adjacent to (**A**) all differentially regulated genes (ALL) or (**B**) upregulated genes (UP), using HOMER known motifs. (related to Figure 5)

**Figure S6. Examples of chromatin accessibility footprint profiles at TF motifs**. (related to Figure 5).

(**A**) Average footprint profiles at transcription factor (TF) binding motifs plotted at base pair resolution, located within accessible chromatin *loci* adjacent to any differentially regulated genes (down or upregulated) for the following TF or binding sites: *Isre*, IRF1, IRF2, NFκB in PBS-treated or IL1B-treated samples, indicating that these TFs might already bind DNA.

(**B**) Average footprints showing sharp internal spikes are suggestive of transposase insertion bias for PU.1-IRF8 or PGR factors, from PBS-treated, IL1B-treated samples or from both conditions (PBS plus IL1B)..

**Figure S7. Constitutive expression of cytokines and chemokines by unstressed early maturating OPCs and immune/inflammatory signature**. (related to Figure 6).

(**A**) ***Expected increase in the expression of genes associated with myelination between P5 and P10 in normal conditions and expected impairment of this increase upon IL1B exposure***. RT-qPCR analyses in P3, P5, and P10 O4+ OPCs. Note that in this set of experiments the downregulation of myelination genes was not observed at P5, a stage at which these genes only start to be expressed.

(**B**) ***O4+ OPCs from naïve pups also constitutively express cytokine and chemokine genes***. RT-qPCR experiments comparing O4+ OPC from naïve (no injection), PBS-injected and IL1B-injected pups, showing that PBS injection does not induce expression of cytokines and chemokine genes, *per se*, since transcripts levels are equivalent in both PBS and naïve conditions, in contrast to what is observed upon IL1B treatment.

(**C**,**D**) **Cross-species comparison confirms global similarities in the chromatin landscapes of HAEC and unstressed O4+ OPC datasets**.

(**C**) Comparison of 100 peaks which were found to be differential in the HAEC (public) dataset upon IL1B treatment, among all (7739) matched peaks between human (HAEC) and mouse (OPCs) datasets. Reads were normalized for each set of peaks against the total number of reads present in all matched peaks and converted into reads per million. Distributions were compared using a one-sample Wilcoxon rank test.

(**D**) Read number distribution of peaks upon IL1B treatment in the HAEC and OPC datasets. Reads were normalized for each set of peaks against the total number of reads present in the 7739 matched peaks and converted into RPM (read per million).

## LIST AND LEGENDS OF SUPPLEMENTAL INFORMATION

### TABLES

**Table S1: Alignment statistics of ATAC-seq**

The alignment statistics of the samples is in line with what is expected from ATAC-Seq samples. Losing in the region of 10% of reads to mitochondrial alignment is normal for this type of data.

**Table S2: Coordinates of the 213**,**246 peaks (mm10) detected in PBS and/or IL1B samples**

MACS2 peak calling was run separately on PBS and IL1B pooled samples (n=3/group). The two resulting peak files (almost 200,000 peaks in each condition) were merged and the mm10 blacklist removed, leading to a list of 213,246 peaks detected in at least one condition (mm10 coordinates).

**Table S3: List and annotation of the 524 differentially accessible peaks**

Reads were counted in each of the 213,246 peaks (Table S2) for each sample individually (3 PBS and 3 IL1B samples). Comparison and statistical analysis with EdgeR (exact test and FDR by Benjamini-Hochberg method) identified 524 peaks with differential accessibility (FDR<0.05). Peaks were annotated using HOMER annotatePeaks.

**Table S4: GO-term Biological Pathway analysis of the 524 differentially accessible peaks**

GO-term Biological Pathway enrichment analysis was performed on the list of gene names (478) associated with the 524 peaks (Table S3) using David6.8.

**Table S5: List of the differentially expressed probes from the microarray analysis**

Agilent microarray data from 6 PBS and 6 IL1B samples (O4+, P5) were submitted to t-test and FDR by Benjamini-Hochberg method. Probes with FDR<0.05 and FC>1.5 (or <-1.5) are listed in the table (Sheet 1). Up-regulated (Sheet 2) and down-regulated (sheet 3) probes are also presented separately. Red flags (columns F and G) indicate the number of undetectable samples (0 means that all samples were detected). Red and green fold change values (column K) correspond to FC>2.0 and FC<-2.0, respectively. Individual values (N to Y columns) are normalized median-centered Log2 intensities.

**Table S6: GO-term Biological Pathway analysis of the UP and DOWN-regulated genes**

GO-term Biological Pathway enrichment analysis was performed using David6.8 on significantly up-regulated (Sheet 1) and down-regulated genes (Sheet 2) from the microarray analysis (Table S5).

**Table S7: Genes whose alterations in expression correlate to opening or closing of the chromatin**

**Table S8: Genes whose alterations in expression correlate to opening or closing of the chromatin in region located +/- 8kb around the TSS**.

**Table S9:** ATAC-Seq peaks associated with upregulated genes and revealing the existence of paired TFBS motifs

**Table S10:** Hogan (HAECS) (list of human gene names and the corresponding orthologue genes in mice, that have been used for the cross-species comparison.

**Table S11: List of the RT-qPCR primers**

**Jupyter notebook: ATAC-seq workflow**

Detailed and explained bioinformatics workflow used for the analysis of ATAC-seq dataset.

## ACKNOWLEDGEMENTS

We are grateful to Dr. Kevin Cheeseman and Dr Magali Hennion (UMR7216) for helpful discussions and comments on the manuscript. This work benefited from equipment and services from the iGenSeq core facility, at the Institut du Cerveau et de la Moëlle (Paris, France). We are particularly grateful to Yannick Marie, the Head of iGenSeq core facility) and Emeline Mundwiller. We are grateful to Dr Sheila Harroch (Pasteur Institute, Paris, France) for the kind gift of the Oli-neu cell line. The authors acknowledge the support of the Freiburg Galaxy Team. We thank the Buffon animal housing facilities at the Jacques Monod Institute and at Robert Debré Hospital (University of Paris, Paris, France).

