## Supplementary figures and images for "Epigenome priming dictates transcription response and white matter fate upon perinatal inflammation"

### Figure S1

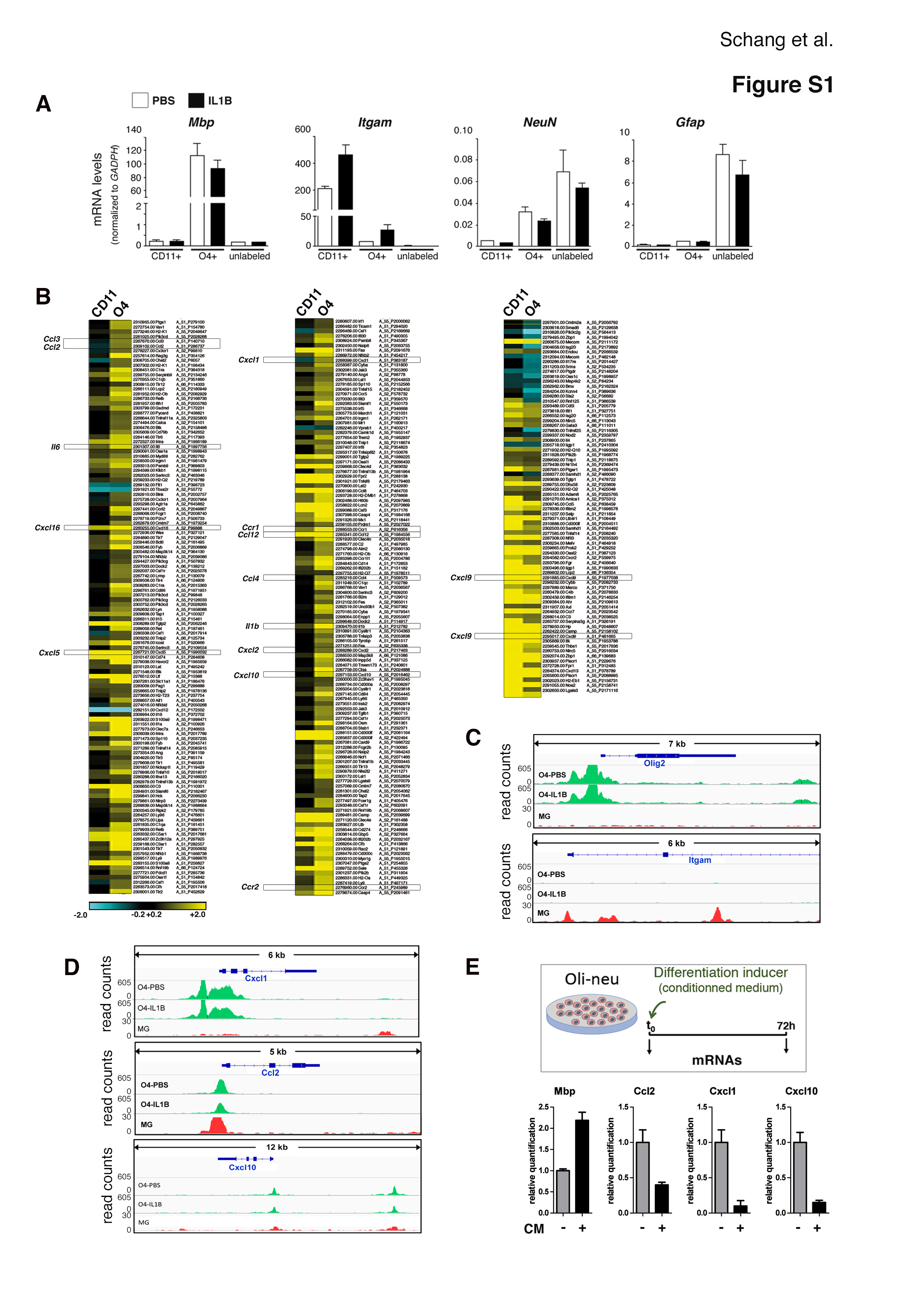

### Figure S2

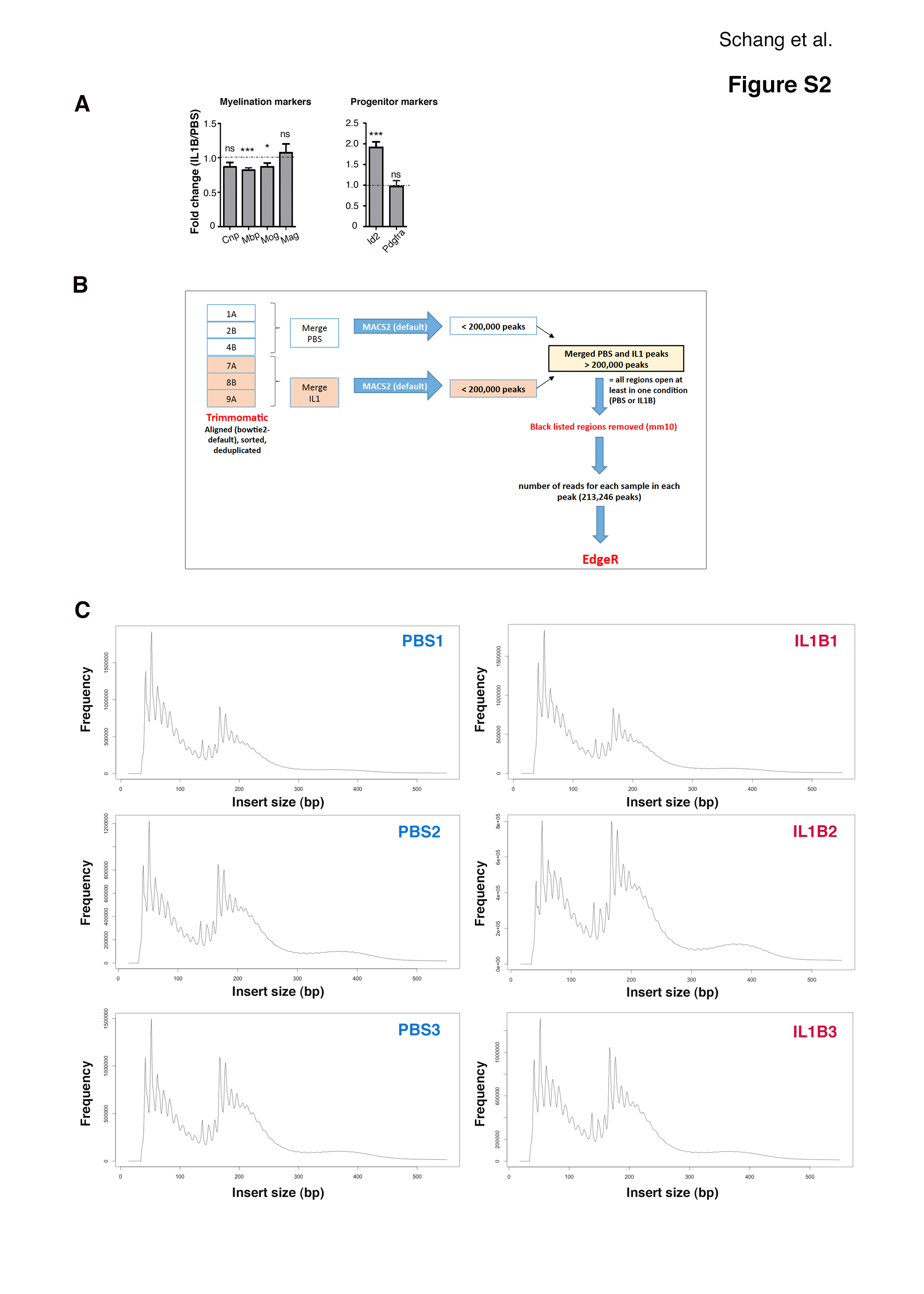

### Figure S3

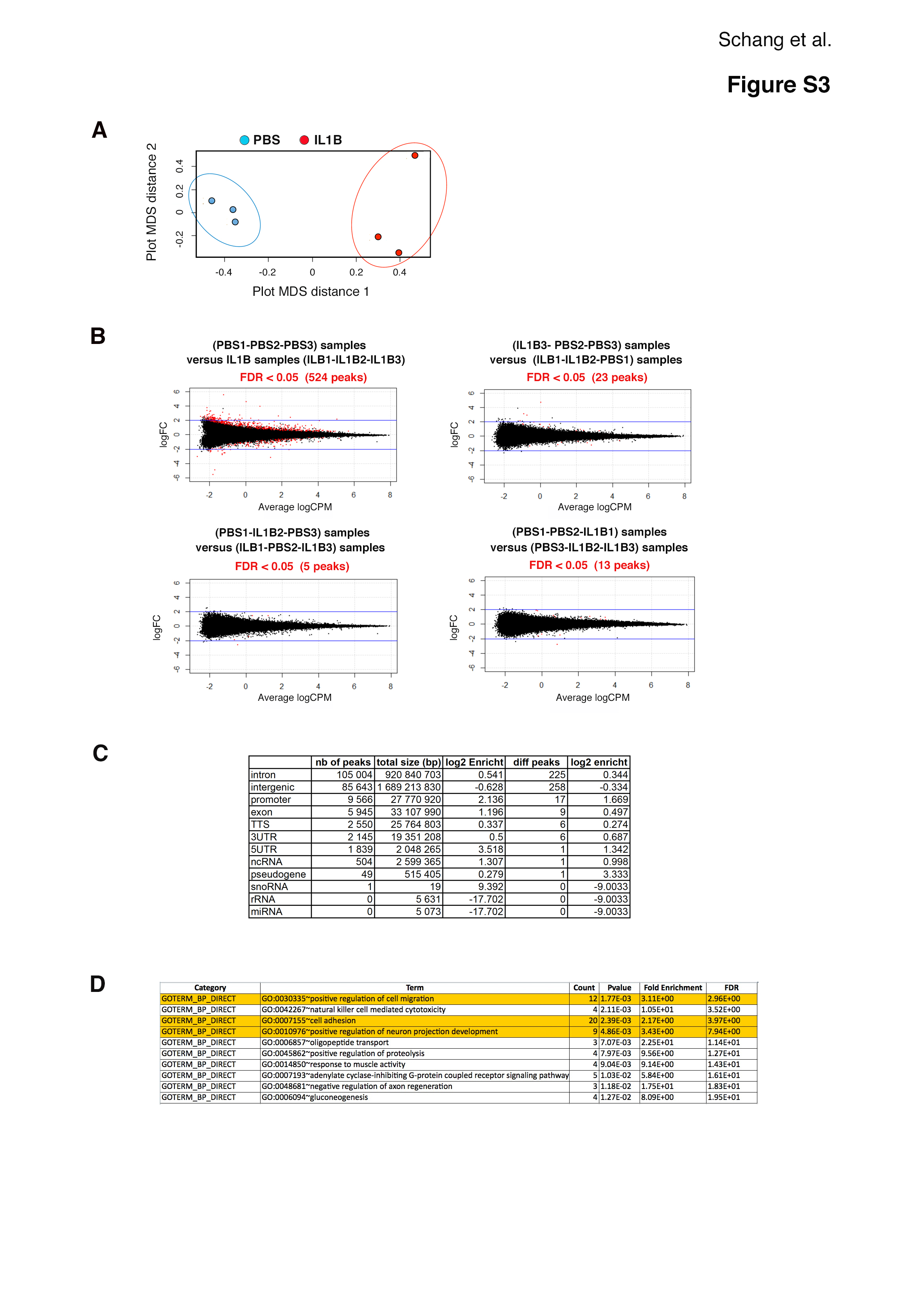

### Figure S4

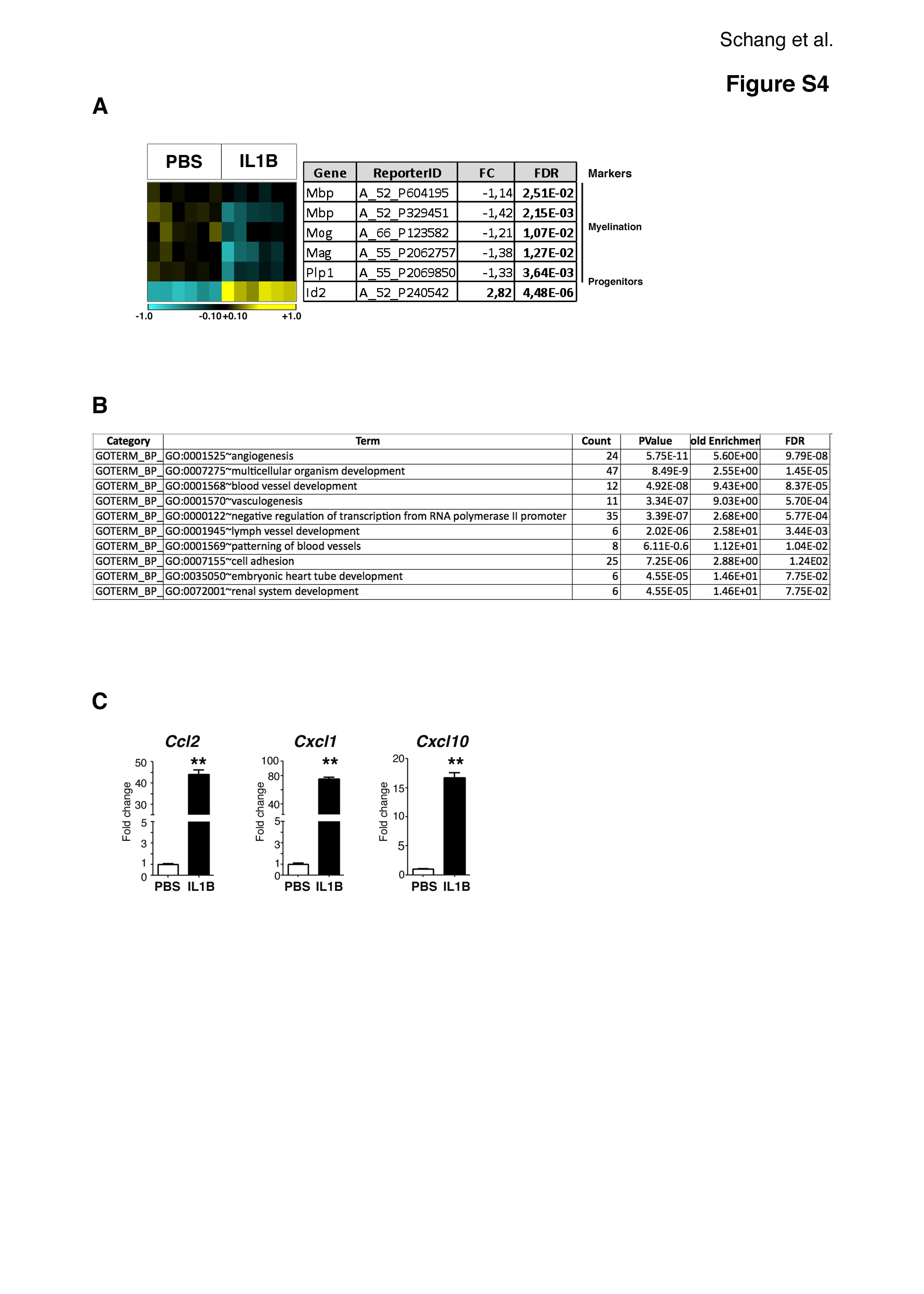

### Figure S5

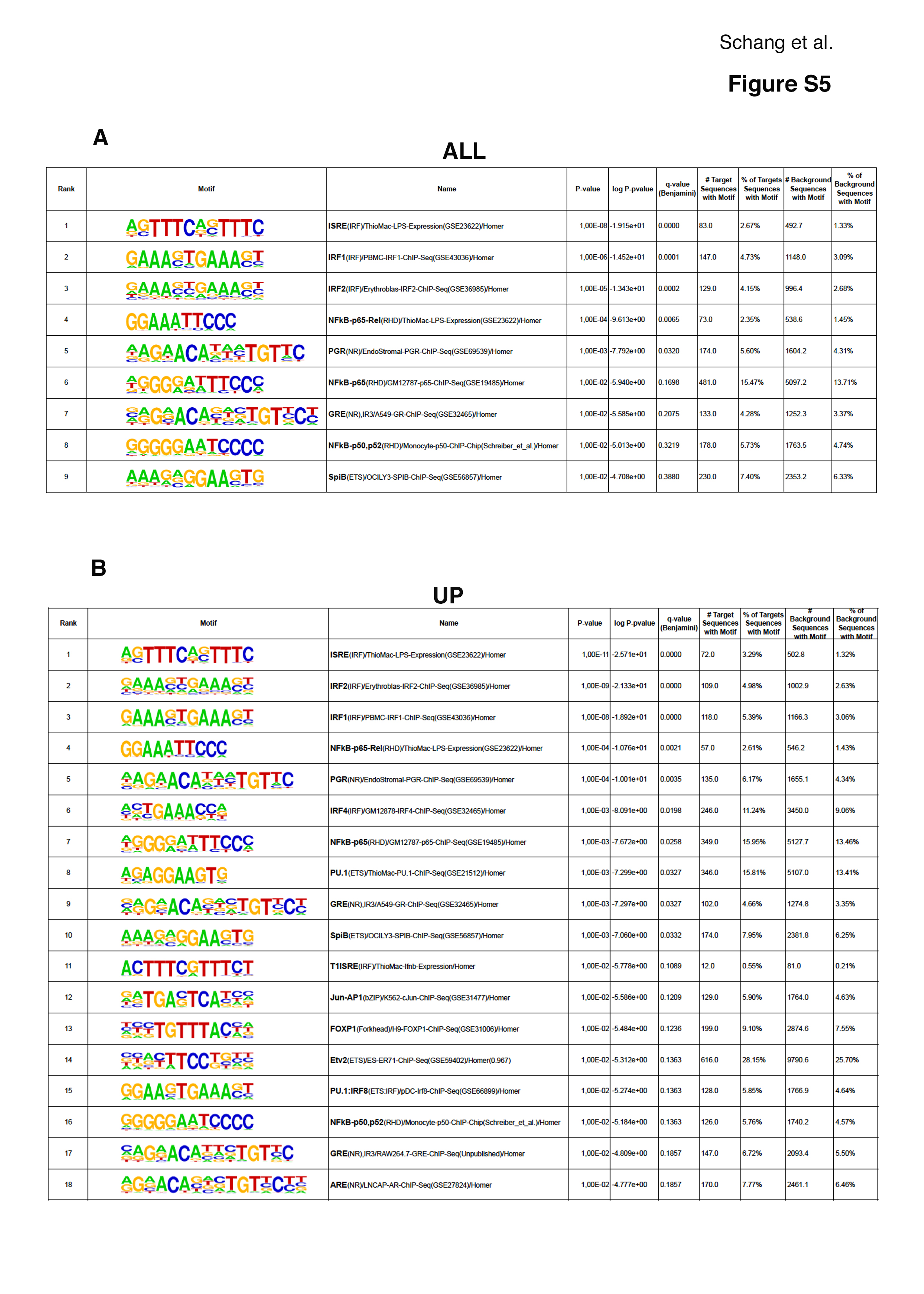

### Figure S6

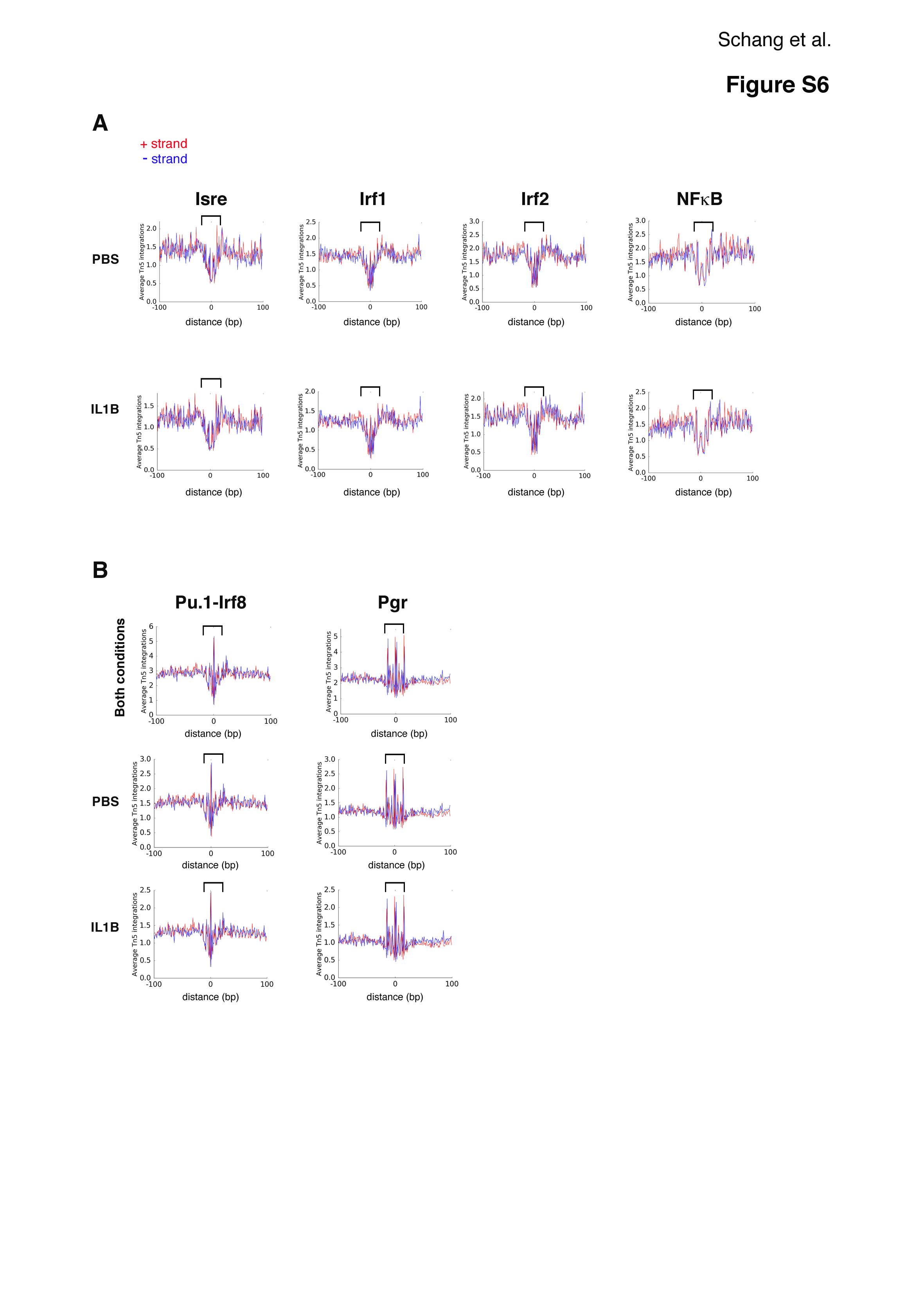

### Figure S7

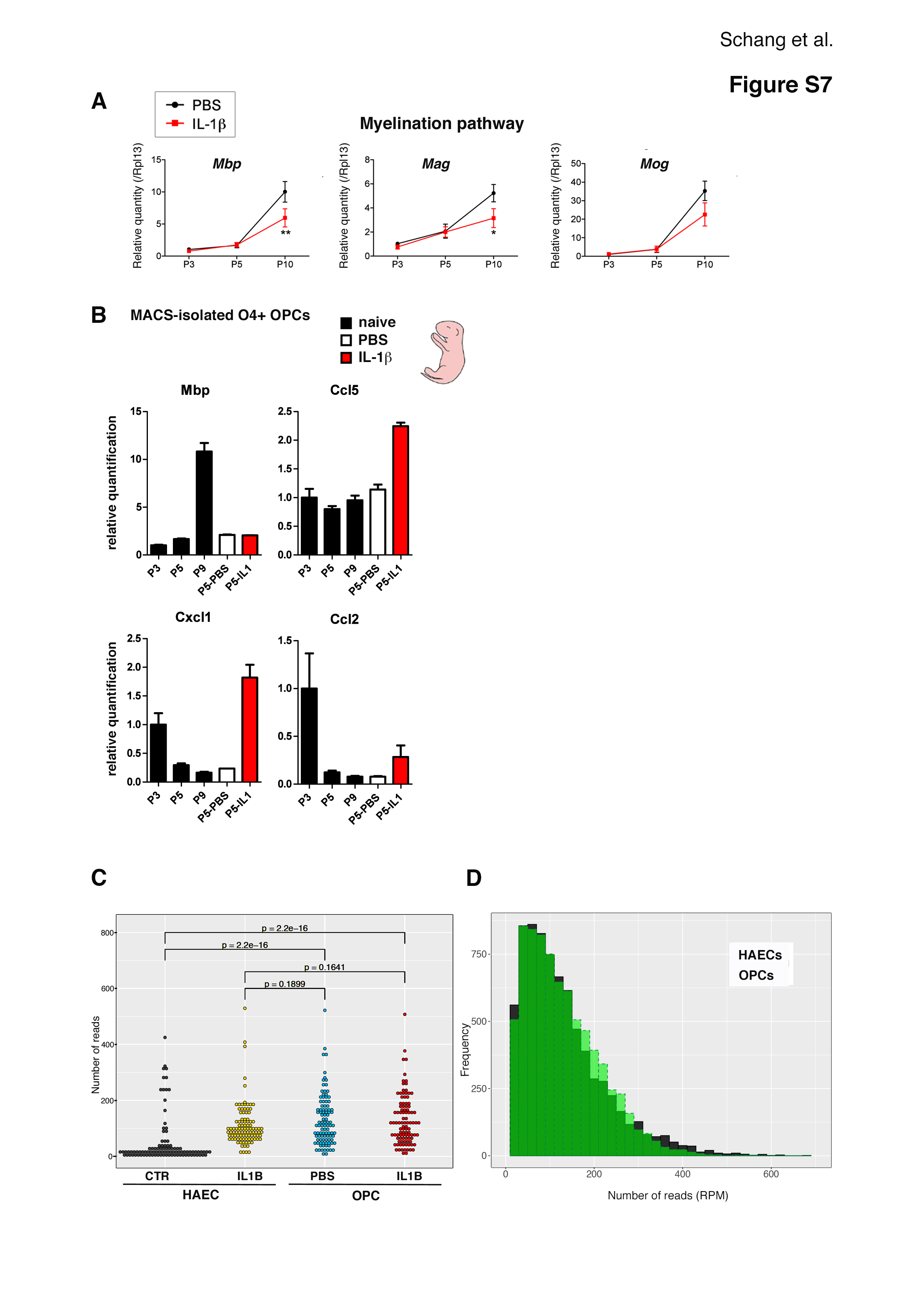
