## Supplementary material for "Epigenome priming dictates transcription response and white matter fate upon perinatal inflammation": Table S1

### Table S1. Alignment Statistics of ATAC-Seq data

| Paired | | PBS1 | PBS2 | PBS3 | IL1B1 | IL1B2 | IL1B3 |
|---|---|---|---|---|---|---|---|
| **Total Reads** | | 73 686 516 | 67 049 336 | 79 305 947 | 74 117 441 | 61 622 256 | 76 360 806 |
| **Removing Mitochondria** | **Remaining Reads** | 66 505 362 | 60 228 153 | 71 141 195 | 66 632 160 | 55 217 126 | 66 962 284 |
| | **% of Total Removed** | 9.75% | 10.17% | 10.30% | 10.10% | 10.39% | 12.31% |
| **% Reads Remaining to Align to Nuclear** | | 90.25% | 89.83% | 89.70% | 89.90% | 89.61% | 87.69% |
| **Nuclear Only** | **Con 1 time only** | 57 751 458 | 52 192 349 | 62 471 914 | 56 958 297 | 47 542 920 | 58 362 195 |
| | **Con >1 time** | 340 054 | 325 951 | 371 952 | 348 211 | 320 270 | 351 148 |
| | **Discon 1 time** | 47 160 | 34 979 | 42 034 | 43 352 | 31 631 | 39 905 |
| | **Without mates 1** | 541 607 | 526 165 | 592 143 | 522 936 | 491 098 | 569 201 |
| | **Without mates >1** | 340 054 | 325 951 | 371 952 | 348 211 | 320 270 | 351 148 |
| | ***Total All Alignments*** | *59 020 333* | *53 405 395* | *63 849 995* | *58 221 007* | *48 706 189* | *59 673 597* |
| | **% of total reads** | 80.10% | 79.65% | 80.51% | 78.55% | 79.04% | 78.15% |

| | | | | | | |
|---|---|---|---|---|---|---|
| **Nuclear Reads Mapping 1 time only** | 58 340 225 | 52 753 493 | 63 106 091 | 57 524 585 | 48 065 649 | 58 971 301 |

| | | | | | | |
|---|---|---|---|---|---|---|
| **% of Total Reads Mapping to nucleus 1 time only** | 79.17% | 78.68% | 79.57% | 77.61% | 78.00% | 77.23% |

The alignment statistics of the samples is in line with what are expected from ATAC-seq samples. Losing in the region of 10% of reads to mitochondrial alignment is usual for this type of data.
